# Transmembrane Serine Protease TMPRSS11B promotes an acidified tumor microenvironment and immune suppression in lung squamous cell carcinoma

**DOI:** 10.1101/2025.04.01.646727

**Authors:** Hari Shankar Sunil, Jean Clemenceau, Isabel Barnfather, Sumanth R. Nakkireddy, Anthony Grichuk, Luke Izzo, Bret M. Evers, Lisa Thomas, Indhumathy Subramaniyan, Li Li, William T. Putnam, Jingfei Zhu, Barrett Updegraff, John D. Minna, Ralph J. DeBerardinis, Jinming Gao, Tae Hyun Hwang, Trudy G. Oliver, Kathryn A. O’Donnell

## Abstract

Lung cancer is the leading cause of cancer-related deaths worldwide. Existing therapeutic options have limited efficacy, particularly for lung squamous cell carcinoma (LUSC), underscoring the critical need for the identification of new therapeutic targets. We previously demonstrated that the Transmembrane Serine Protease *TMPRSS11B* promotes transformation of human bronchial epithelial cells and enhances lactate export from LUSC cells. To determine the impact of TMPRSS11B activity on the host immune system and the tumor microenvironment (TME), we evaluated the effect of *Tmprss11b* depletion in a syngeneic mouse model. *Tmprss11b* depletion significantly reduced tumor burden in immunocompetent mice and triggered an infiltration of immune cells. RNA FISH analysis and spatial transcriptomics in the autochthonous *Rosa26-Sox2-Ires-Gfp^LSL/LSL^*; Nkx2-1^fl/fl^; *Lkb*1^fl/fl^ (SNL) model revealed an enrichment of *Tmprss11b* expression in LUSC tumors, specifically in Krt13^+^ hillock-like cells. Ultra-pH sensitive nanoparticle imaging and metabolite analysis identified regions of acidification, elevated lactate, and enrichment of M2-like macrophages in LUSC tumors. These results demonstrate that TMPRSS11B promotes an acidified and immunosuppressive TME and nominate this enzyme as a therapeutic target in LUSC.

## Introduction

Lung cancer is the leading cause of cancer related deaths worldwide ^1^. Lung cancer is broadly classified into small cell (SCLC) and non-small cell lung cancer (NSCLC), with the latter representing ∼85% of all lung cancers. Lung squamous cell carcinoma (LUSC) is one type of NSCLC, accounting for ∼30% of all lung cancer cases ^2,3^. These tumors are highly heterogenous and lack targeted therapies. Immune checkpoint inhibitors have emerged as the first line therapy for LUSC patients, but this approach is effective only in a subset of patients ^4–8^. This highlights a critical need for the identification and characterization of new therapeutic targets in LUSC.

Basal cells are thought to be one cell of origin for LUSCs, although accumulating evidence suggests that LUSCs may arise from multiple cell types including club cells and alveolar type II cells ^9–15^. Hillock cells are a recently described cell type in the lung that were shown to be an injury-resistant reservoir of stem-like cells. They represent a distinctive population of basal stem cells that express Keratin13 (KRT13) and other genes associated with barrier function, cell adhesion, and immunomodulation. Interestingly, hillock cells were also shown to be one origin of squamous metaplasia, a precursor to LUSC ^16–19^. A better understanding of hillock cell biology and the origins of LUSC may lead to new therapies for this tumor type.

With the goal of discovering novel genes that drive lung tumorigenesis, we previously performed an unbiased *Sleeping Beauty* (SB)-mediated transposon mutagenesis screen in immortalized human bronchial epithelial cells (HBECs). This screen revealed that the transmembrane serine protease TMPRSS11B promotes the transformation of HBECs and enhances LUSC tumor growth in immunocompromised mice ^20^. TMPRSS11B belongs to the <u>d</u>ifferentially <u>e</u>xpressed in <u>s</u>quamous cell <u>c</u>ancer (DESC) family of genomically clustered, trypsin-like serine proteases that share commonality in their type-II transmembrane insertion, catalytic triad spacing, and disulfide-bonding anchoring their serine protease domain to a membrane proximal cysteine ^21^. We found that *TMPRSS11B* expression is highly upregulated in LUSCs compared to normal lung tissues ^22^. Mechanistic studies and metabolomics further revealed that TMPRSS11B interacts with the lactate monocarboxylate transporter 4 (MCT4) and its obligate chaperone, Basigin (CD147). TMPRSS11B catalytic activity promotes solubilization of Basigin, which enhances MCT4-mediated lactate export, glycolytic flux, and tumor growth, thereby promoting tumorigenesis in LUSC ^20^.

Although lactate is usually generated as a by-product of enhanced glycolytic flux in tumor cells, there is significant heterogeneity in its metabolism. Some tumor types export of lactate through MCT4 in order to increase glycolysis to drive tumorigenesis. In contrast, other tumor types employ monocarboxylate transporter 1 (MCT1) to import lactate, which can be used for energy production during tumorigenesis ^23–25^. In addition, extracellular lactate is reported to influence the tumor microenvironment to support tumor growth by inhibiting T-cell function ^26,27^, recruiting Tregs ^28^, inducing PD-L1 expression ^29^, and by polarizing macrophages to the M2 or immunosuppressive subtype ^30–33^. These studies suggest that lactate metabolism is a vulnerability that may be harnessed for therapeutic targeting of cancer cells.

Tumor-associated macrophages (TAMs) represent a significant fraction of immune cells in the tumor microenvironment and have been shown to promote tumor progression by promoting epithelial-to-mesenchymal transition (EMT), extracellular matrix remodeling through the secretion of proteolytic enzymes, exhaustion and suppression of cytotoxic T-cells, and recruitment of Tregs ^34–38^. The M2 subtype is immunosuppressive and tumor promoting, while the M1 subtype is anti-tumorigenic. In NSCLC, largely LUSCs, M2 macrophages represent a substantial proportion of TAMs, with increased enrichment in the tumor stroma. Moreover, higher M1/M2 ratios correlate with better survival for patients, demonstrating the impact of immunosuppressive TAM subtypes in lung cancer ^39–42^.

Using an autochthonous model of LUSC coupled with spatial transcriptomics, ultra pH sensitive nanoparticle imaging and metabolomics, we show that *Tmprss11b*-expressing lung squamous tumors and the surrounding microenvironment exhibit elevated lactate levels and accumulate immunosuppressive M2-like tumor associated macrophages. We also demonstrate that *Tmprss11b* is restricted to squamous tumors and is enriched specifically in Krt13^+^ hillock like cells. Collectively, these studies reveal the establishment of an acidified and immunosuppressive TME in *Tmprss11b*-high LUSC and suggest a new therapeutic approach of targeting this enzyme in the hillock-like population of squamous lung cancer.

## Results

### *Tmprss11b* depletion reduces tumor growth and enhances CD4+ T cell infiltration

We previously demonstrated that TMPRSS11B inhibition limits tumor growth of human LUSCs in xenograft assays in immunocompromised NOD/SCID Il2rψ**^-/-^** (NSG) mice. However, these studies did not fully recapitulate all aspects of tumorigenesis such as contributions from the tumor microenvironment and the immune system. Based on our previous demonstration that TMPRSS11B promotes lactate export ^20^ and the role of lactate in modulating immune cell function, we hypothesized that loss of function of TMPRSS11B would reduce tumor growth and enhance immune cell infiltration in the TME in immunocompetent mice. To test this hypothesis, we used CRISPR/Cas9 to knockout *Tmprss11b* in KLN205 cells, an established syngeneic mouse model of murine lung squamous cell carcinoma (**Fig. 1A-B, Supplementary Fig. S1A**) ^43^. Cells were transplanted subcutaneously in immunocompetent DBA/2 WT mice and tumor volumes were assessed. Tumor growth was significantly reduced in *Tmprss11b* knockout tumors compared to control tumors (**Fig. 1C, Supplementary Fig. S1B**). We also used doxycycline-inducible short hairpin RNAs (shRNAs) to knockdown *Tmprss11b* in KLN205 cells after tumor initiation (**Fig. 1D**). Cells were transplanted into immunocompetent DBA/2 mice, and mice were maintained on doxycycline water to induce shRNA expression after tumor formation. We observed strong impairment of tumor growth following depletion of *Tmprss11b* (**Fig. 1E, Supplementary Fig. S1C-D**). Efficient knockdown of *Tmprss11b* was confirmed in *Tmprss11b*-shRNA expressing tumors (**Supplementary Fig. S1E**). Fluorescent immunohistochemistry (IHC-F) demonstrated that *Tmprss11b* loss of function triggered an accumulation of CD4+ T cells (**Fig. 1F**), and to a lesser extent CD8+ T cells (**Supplementary Fig. S1F**), in syngeneic tumors. Moreover, we observed a significant decrease in phospho-ERK signaling (**Fig. 1G**). These findings establish that *Tmprss11b* loss of function alters immune cell infiltration in the TME and suppresses the MAPK signaling pathway.

**Figure 1.**
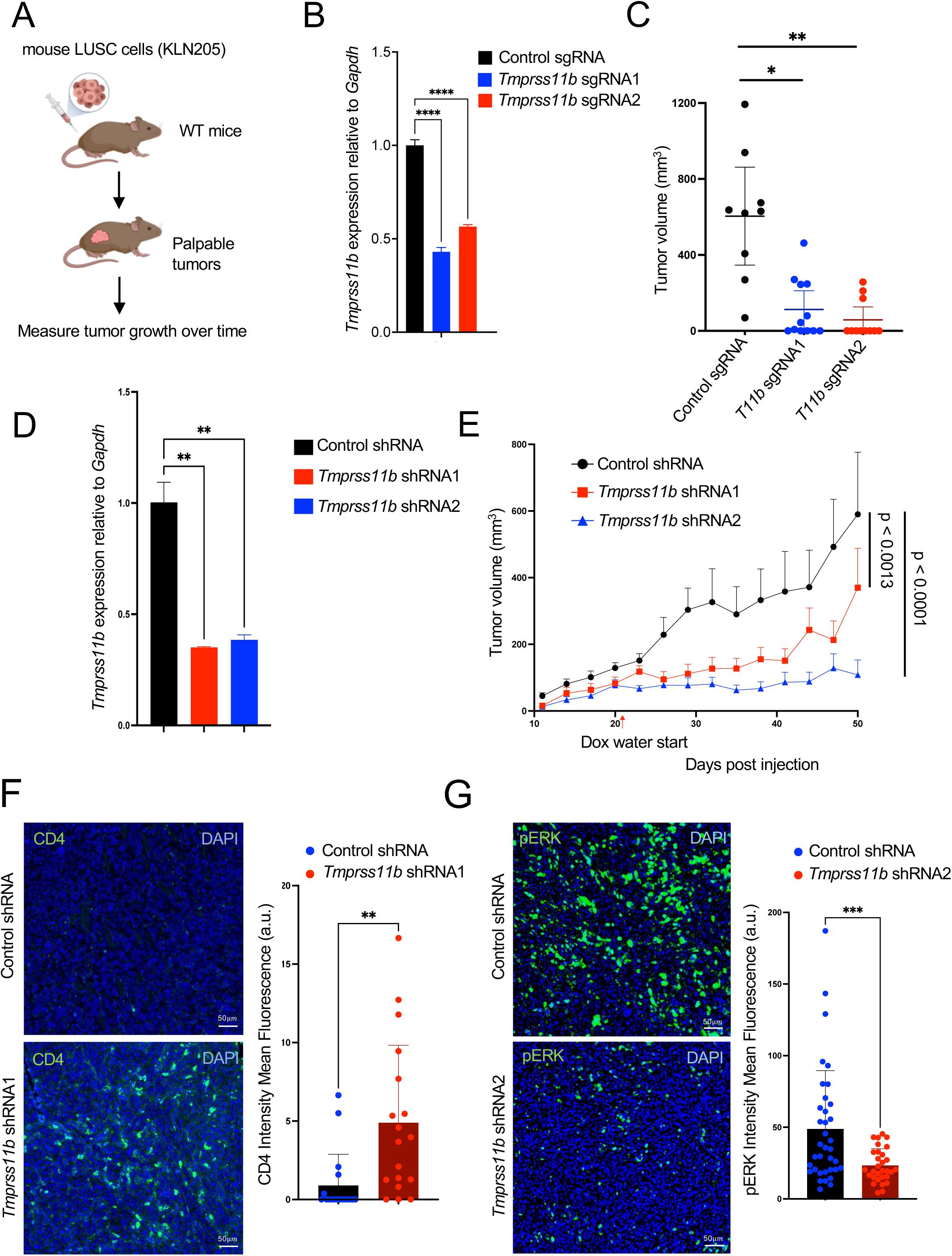
*Tmprss11b* depletion reduces tumor growth and enhances immune cell infiltration. **A)** Schematic of the syngeneic experiments performed using KLN205 murine LUSC cells. **B)** qRT-PCR analysis of *Tmprss11b* mRNA in KLN205 cells expressing control sgRNA or two independent *Tmprss11b* sgRNA. **C)** Quantification of tumor volumes of KLN205 tumors expressing control or *Tmprss11b* sgRNA on day 45 (terminal measurement) post injection in syngeneic DBA/2 wild-type mice (n=9 control sgRNA mice; n=12 *Tmprss11b* sg1 mice; n=12 *Tmprss11b* sg2 mice). Brown-Forsythe and Welch ANOVA test with Dunnett’s T3 multiple comparisons test was used for the statistical analysis. **D)** qRT-PCR analysis of *Tmprss11b* mRNA in KLN205 cells expressing doxycycline-inducible control shRNA or two independent shRNA sequences targeting *Tmprss11b*. **E)** Quantification of tumor volumes of KLN205 cells expressing doxycycline-inducible control or *Tmprss11b* shRNA in syngeneic DBA/2 wild-type mice. (n=9 control shRNA mice; n=6 *Tmprss11b* sh1 mice; n=8 *Tmprss11b* sh2 mice). Linear mixed-effects models were used to investigate the differences in tumor volume over time among the three groups. **F)** Fluorescent immunohistochemistry (IHC-F) staining for CD4+ T cells in KLN205 tumor sections, with quantification. An unpaired t test with Welch’s correction was used to compare CD4+ T cells between the groups (n=10 fields per tumor section, 3 tumors per group). **G)** Fluorescent immunohistochemistry (IHC-F) staining for phosphorylated ERK (pERK) in KLN205 tumors, with quantification (right). An unpaired t test with Welch’s correction was used to compare phosphorylated ERK levels between the groups (n=10-14 fields per tumor section, 3 tumors per group).

### *Tmprss11b* expression is enriched in lung squamous tumors

We next investigated the expression of *Tmprss11b* in a panel of tumors derived from autochthonous mouse models of lung cancer representing lung adenocarcinoma (LUAD), lung squamous (LUSC), and small cell lung cancer (SCLC). *Tmprss11b* is highly expressed in LUSC tumors, with the highest expression in the *Rosa26-Sox2-Ires-Gfp^LSL/LSL^*;*Nkx2-1^fl/fl^*;*Lkb1^fl/fl^* (SNL) mouse model (**Fig. 2A**). This model, based on overexpression of the *Sox2* transcription factor and loss of the tumor suppressors *Lkb1* and *Nkx2-1*, accurately recapitulates the histology and microenvironment of LUSCs ^44^. Interestingly, these mice develop LUSC and mucinous LUAD, thereby providing an ideal setting to investigate intra-tumoral heterogeneity and metabolism.

**Figure 2.**
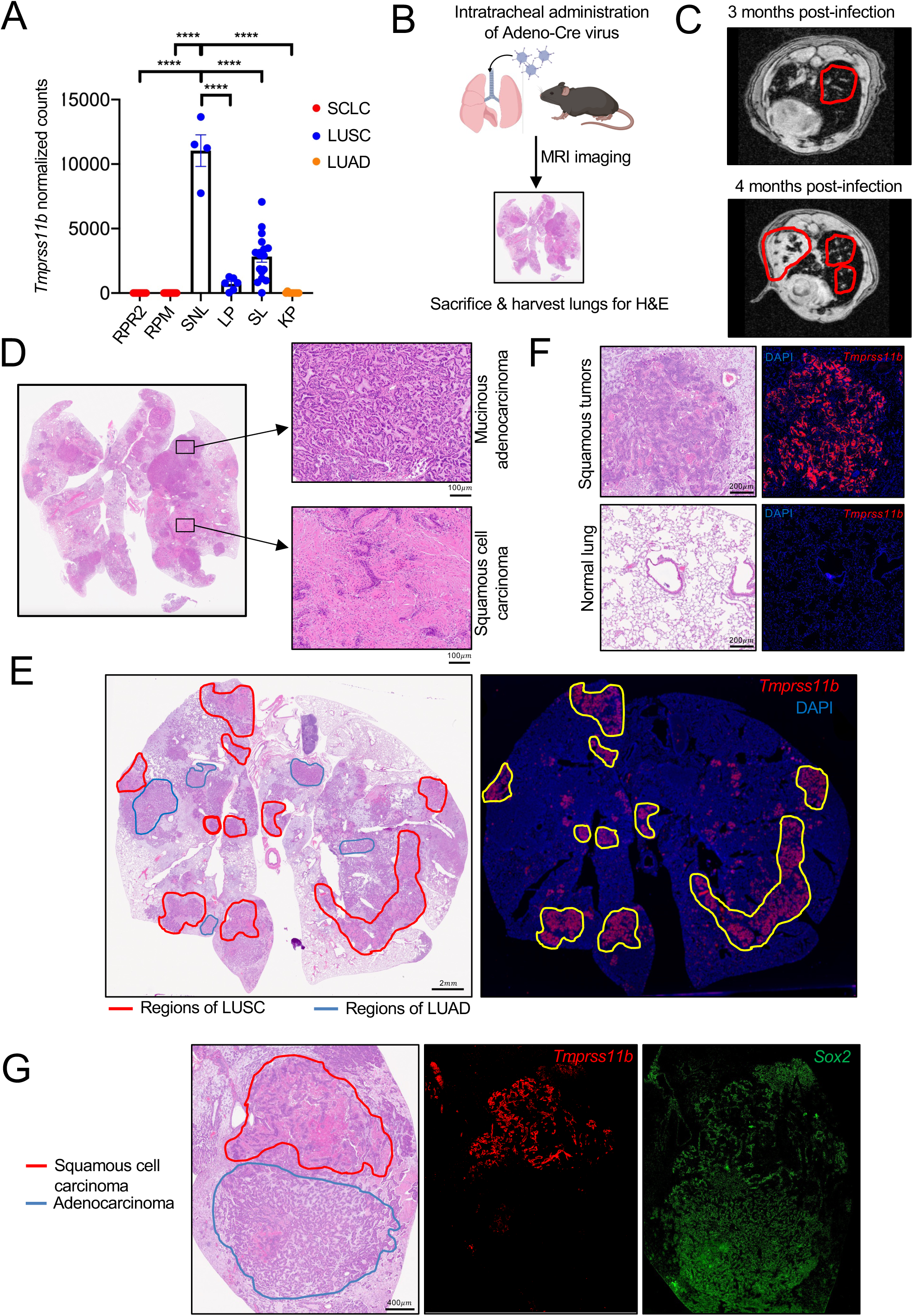
*Tmprss11b* expression is enriched in mouse lung squamous tumors. **A)** RNA sequencing analysis of tumors from various mouse models of lung cancer. The y-axis represents the normalized counts for *Tmprss11b*. **B)** Schematic representation of Ad-Cre mediated tumor induction in SNL mice. **C)** Representative MRI images of the mice in B), 3-& 4-months post infection. Red outlines denote tumors. **D)** Representative H&E images of SNL mouse lung, 7 months post infection with Ad-Cre showing distinct regions of LUSC and mucinous LUAD. **E)** H&E image of SNL mouse lung (11 months post infection with Ad-Cre) and RNAscope of *Tmprss11b* on a serial section. Left, red outline denotes squamous tumors based on H&E staining. Right, yellow outline denotes regions with *Tmprss11b* expression (red) corresponding to the regions of squamous tumors. **F)** Zoom-in of (E) showing *Tmprss11b* expression by RNAScope in squamous tumors (top panel) and normal lung (bottom panel). **G)** Representative H&E image with annotations and RNAscope analysis of *Tmprss11b* (red) and *Sox2* (green) in SNL lung sections.

We initiated tumorigenesis in SNL mice through intratracheal administration of adenovirus expressing Cre (Ad-Cre) (**Fig. 2B**) and monitored tumor progression over time (**Fig. 2C**). At 8-months post-infection, we observed appreciable lung tumor burden and distinct regions of squamous tumors (LUSC) and mucinous adenocarcinomas (LUAD) identified by H&E staining (**Fig. 2D**). We next assessed the spatial distribution of *Tmprss11b* expression with RNA fluorescent *in situ* hybridization (FISH) (RNAscope). We observed selective enrichment of *Tmprss11b* in squamous lung tumors but not in mucinous adenocarcinomas or normal lungs (**Fig. 2E-G**). In contrast, *Sox2*, the oncogenic driver in this model, was expressed in LUSC and LUAD tumors (**Fig. 2G**). These data demonstrate that *Tmprss11b* is selectively expressed in murine lung squamous tumors.

### *Tmprss11b* expression correlates with an increase in squamous markers and oncogenic signaling pathways

Given the heterogeneity observed in SNL lung tumors, we next investigated the gene expression pathways that associate with *Tmprss11b* expression using spatial transcriptomics. High quality sequencing reads were obtained with the Visium CytAssist (10x Genomics) platform (**Supplementary Fig. S2A**) and analysis revealed multiple clusters, indicating spatial heterogeneity (**Fig. 3A**). We annotated the sections as LUSC and LUAD based on histology for additional analyses, including 3 regions as LUSC (S1, S2, S3) and 2 regions as LUAD (A1, A2) (**Fig. 3B**). We assessed the distribution of *Tmprss11b* transcripts and observed unique enrichment in the LUSCs, validating our RNA-FISH results (**Fig. 3C**). As expected, the squamous marker *Trp63* and the mucinous adenocarcinoma marker *Hnf4a* exhibited enrichment in LUSCs and LUADs, respectively (**Fig. 3C**).

**Figure 3.**
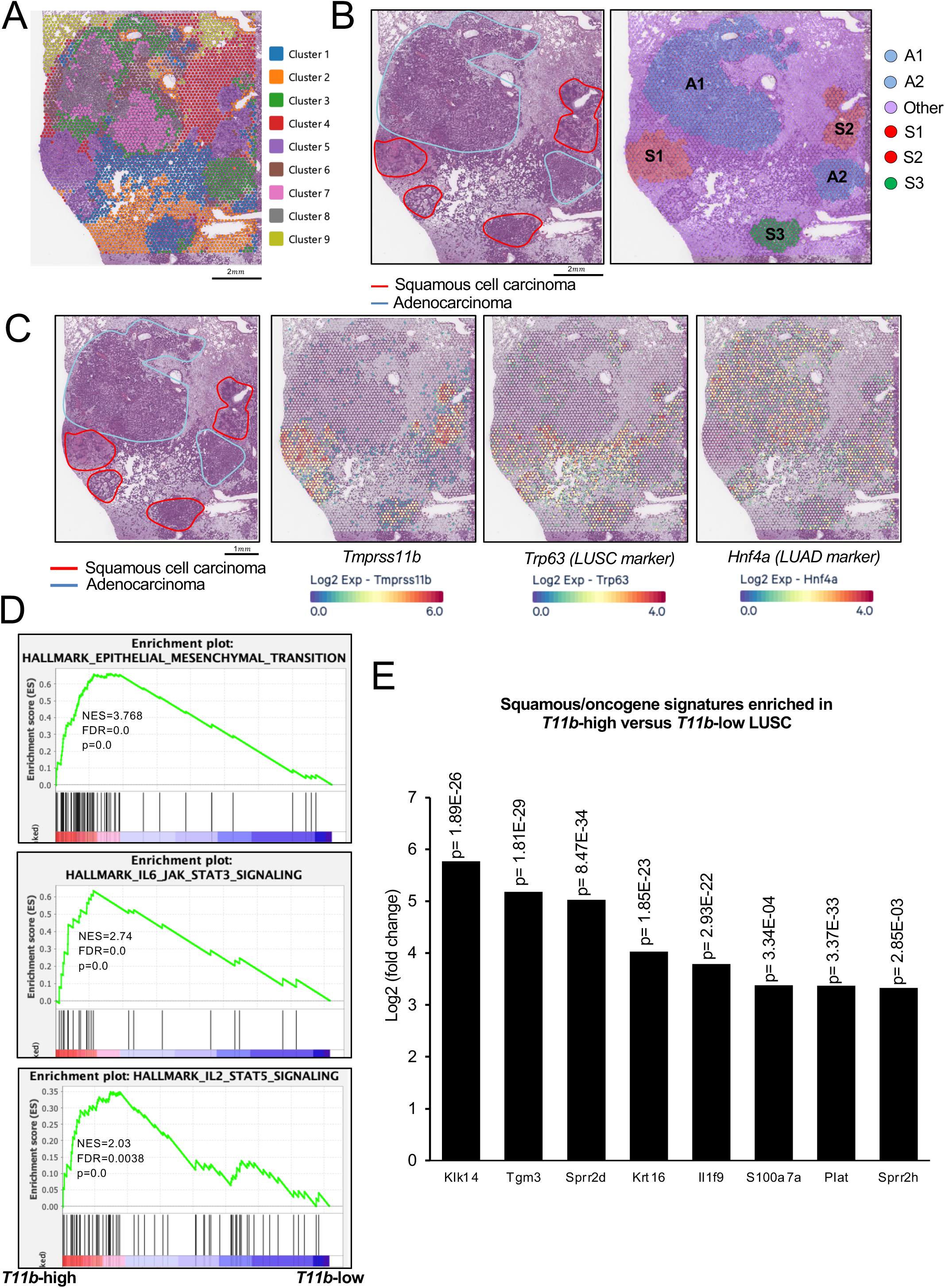
*Tmprss11b* expression correlates with an increase in squamous markers and oncogenic signaling pathways. **A)** Image depicting the different clusters identified in a SNL lung section based on spatial transcriptomic analysis. **B)** H&E image of the lung section from A) (left) and manual annotation of different regions (right); LUSC (S1, S2, S3) and LUAD (A1 and A2). **C)** H&E image and corresponding spatial plots depicting the distribution of the indicated mRNAs. **D)** Gene set enrichment analysis (GSEA) from *T11b*-high versus low squamous tumors (S1&S2 versus S3) with normalized enrichment scores (NES), false discovery rate (FDR) and p values for the indicated gene signatures. **E)** Bar graph representing top squamous genes/oncogenes from differential gene expression (DEG) analysis of *Tmprss11b*-high versus low LUSC spatial transcriptomics data.

We next quantified *Tmprss11b* expression in all annotated regions. Although all LUSC regions expressed higher levels of *Tmprss11b* compared to LUAD regions, the S1 and S2 regions exhibited ∼4-fold higher levels of expression compared to the S3 region (**Supplementary Fig. S2B-C**). To elucidate pathways elevated in squamous tumors with high *Tmprss11b* expression, we performed gene set enrichment analysis (GSEA) on *Tmprss11b-*high (S1, S2) versus *Tmprss11b*-low (S3) LUSC regions and observed an enrichment of oncogenic and immune cell signaling pathways (**Fig. 3D**). In addition, we performed differential gene expression (DEG) analysis on *Tmprss11b-*high versus low regions and observed an increase in genes known to be elevated in squamous tumors (*Krt16* and *Sprr2d*) and genes that promote tumorigenesis, including *Tgm3*, *Krt16*, and *S100a7a* (**Fig. 3E**) ^12,45–47^. We also performed GSEA and DEG comparing *Tmprss11b*-high LUSCs (S1, S2) versus LUADs (A1, A2) and observed a similar enrichment of genes and pathways (**Supplementary Fig. S2D-E**). Immunohistochemistry (IHC) analysis of the LUSC markers KRT16 and LYPD3 demonstrated increased expression in LUSCs compared to LUADs, validating our spatial transcriptomics data (**Supplementary Fig. S2F**). These findings demonstrate that tumors expressing high levels of *Tmprss11b* exhibit an enrichment for squamous markers and an oncogenic gene expression signature.

### TMPRSS11B is enriched in the hillock cell population and induced by KLF4

Given the robust expression of *Tmprss11b* in squamous tumors and the strong correlation with squamous markers and oncogenic signaling pathways, we hypothesized that TMPRSS11B may be expressed in the cell of origin in LUSCs. Basal cells are hypothesized to be the precursor of squamous tumors ^9,10^. However, lineage tracing studies have demonstrated that squamous tumors can arise from multiple cell types in the lung, including club cells and alveolar type II (AT2) cells ^11–15^. Recent findings have identified a new cell type in the pseudostratified lung epithelium known as hillock cells that can give rise to squamous metaplasia, a precursor of squamous tumors ^16^. These are collections of multilayered injury resistant cells that express KRT13 and consist of keratinized upper layers of luminal cells that protect underlying layers of KRT13^+^, TP63^+^ basal stem cells ^16–19,48^. Interestingly, these domains harbor a unique population of basal stem cells that express genes associated with barrier function, cell adhesion, and immunomodulation ^17,48–53^. Moreover, there is a growing appreciation that membrane-anchored serine proteases are important regulators of epithelial development and barrier function ^54^. RNA sequencing of mouse syngeneic tumors revealed that keratin genes were downregulated in *Tmprss11b* depleted cells (**Supplementary** Fig. 3A). Conversely, DEG analysis of *Tmprss11b*-high squamous tumors and LUSC patient data from TCGA uncovered an enrichment of several keratin genes including *KRT13, KRT6, KRT10,* and *KRT14* (**Supplementary** Fig. 3B-E). Given these observations, we sought to determine if TMPRSS11B is expressed in KRT13^+^ hillock cells.

Findings presented in a companion study (Izzo et al. 2025, co-submitted) identified a hillock cell population in human and mouse lung squamous cell carcinomas. Consistent with this observation, we find that *Tmprss11b* is co-expressed with *Keratin 13* (*Krt13)* and *Keratin 6a* (*Krt6a*) in mouse LUSC tumors (**Fig. 4A**). Using hillock, basal and mucinous gene signatures, we assigned regions of enrichment for these signatures in our spatial transcriptomics dataset. We find that the hillock and basal gene signatures were highly enriched in the *Tmprss11b*-high squamous tumors, while the mucinous gene signature was expressed in the adenocarcinomas (**Supplementary** Fig. 4A**-B**). We performed RNA-FISH analysis of *Tmprss11b*, *Trp63* and *Krt13* in SNL tumors and observed an enrichment of *Trp63* in the basal layer and expression of *Krt13* in the suprabasal layer in squamous tumors, consistent with a recent study ^16^ (**Fig. 4B**). Interestingly, *Tmprss11b* expression was suprabasal to *Trp63* and co-expressed with *Krt13*.

**Figure 4.**
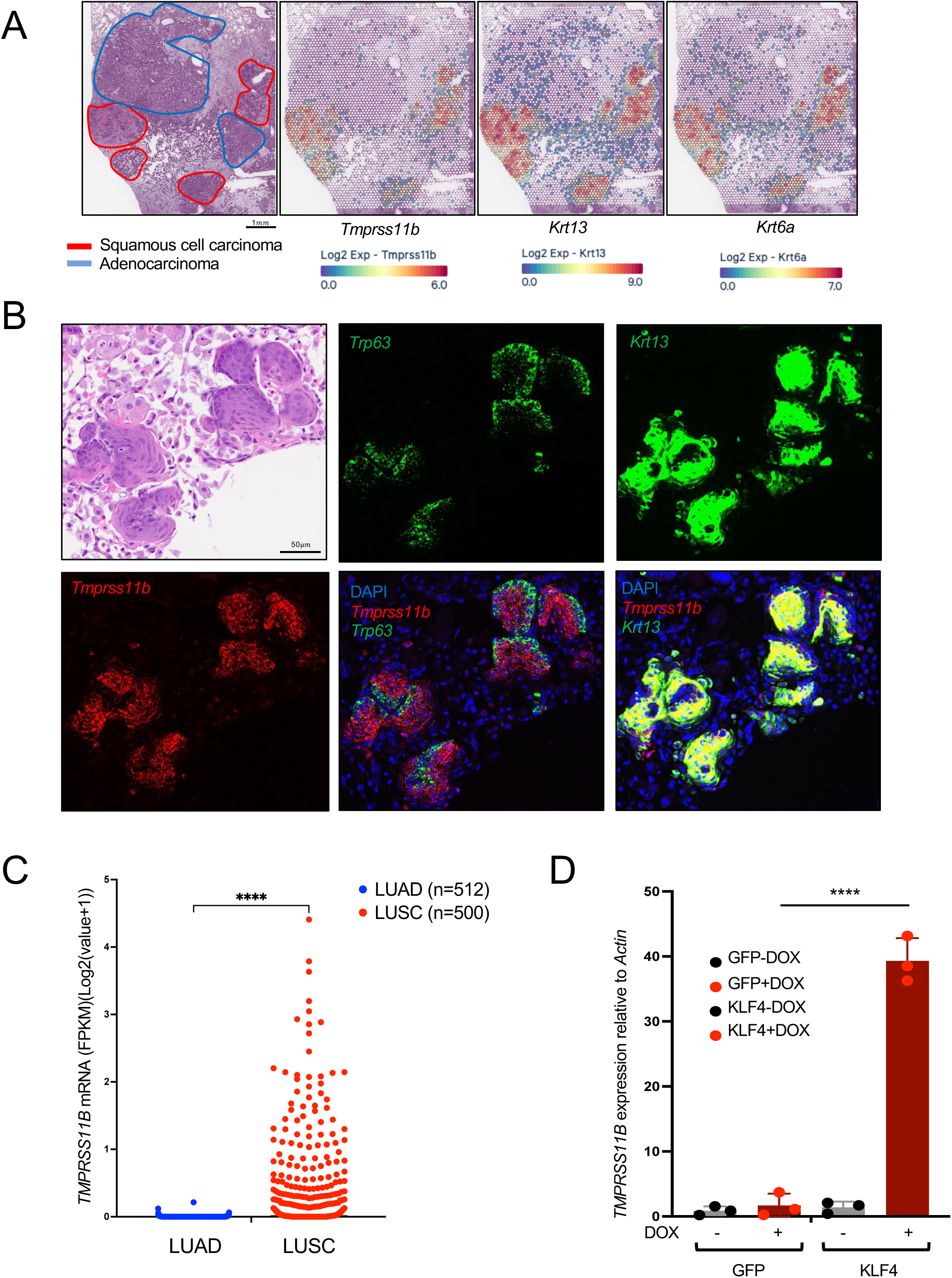
TMPRSS11B is enriched in the KRT13^+^ hillock like cells and induced by KLF4. **A)** H&E image (left) and corresponding spatial plots depicting the distribution of the indicated mRNAs. **B)** Representative H&E image and RNAscope analysis of *Tmprss11b*, *Trp63, Krt13*, and *Krt6a* in SNL lung sections. **C)** *Tmprss11b* transcript abundance in LUSC patient tumors relative to LUAD tumors (data obtained from TCGA-GDC). **D)** qRT-PCR analysis of *Tmprss11b* mRNA in BEAS-2B cells expressing doxycycline-inducible GFP or KLF4.

To extend these findings to human lung squamous tumors, we analyzed LUSC RNA sequencing data from TCGA. Indeed, TMPRSS11B was upregulated in human LUSCs compared to LUAD (**Fig. 4C**). Interestingly, a detailed analysis of cell clusters using a hillock gene signature found that a *TMPRSS11B*-high cluster also expresses *KRT13* (Izzo et al. 2025). Importantly, this study demonstrated that the Krüppel-like factor 4 (KLF4) transcription factor induces a hillock-like gene signature in human bronchial epithelial cells (BEAS-2B). Given the importance KLF4 in regulating a hillock cell state and its regulatory role in a variety of cancers ^55–58^, we sought to determine if TMPRSS11B is induced by KLF4. We performed quantitative RT-PCR for *TMPRSS11B* on RNA isolated from BEAS-2B cells following doxycycline inducible expression of green fluorescent protein (GFP) or KLF4. Indeed, *TMPRSS11B* was induced by ∼40-fold in KLF4-expressing BEAS-2B cells (**Fig. 4D**). Overall, these data show that *TMPRSS11B* is induced by KLF4 and is expressed in the KRT13+ hillock cell population in lung squamous tumors.

### *Tmprss11b*-high squamous tumors exhibit infiltration of M2-like macrophages

Given the enrichment of squamous markers and oncogenic signaling in *Tmprss11b*-expressing squamous tumors (**Fig. 3D**, **Supplementary** Fig. 2D), and the known link between lactate metabolism and immune regulation, we next examined the immune cell populations that infiltrate LUSC tumors. Known markers of neutrophil infiltration including *Cd11b* (*Itgam*), *Cxcl3* and *Cxcl5* were expressed in LUSCs, consistent with prior studies in this mouse model (**Supplementary** Fig. 5A) ^44^. We also performed cell deconvolution analysis and identified an enrichment of additional immune cells subsets, including classical monocytes, alveolar macrophages and leukocytes, with high expression of *Tmprss11b* (**Fig. 5A-B, Supplementary** Fig. 5B-C). GSEA of *Tmprss11b*-high (S1, S2) vs. *Tmprss11b*-low LUSC (S3) and *Tmprss11b*-high LUSC (S1, S2) vs. LUAD (A1, A2) revealed an enrichment for similar pathways (**Fig. 5C Supplementary** Fig. 5D). Given the presence of opposing subtypes of macrophages observed in the tumor microenvironment, based on their ability to promote or suppress tumor progression, we assessed macrophage subtypes that were enriched with higher levels of *Tmprss11b*. DEG analysis of the spatial data identified an enrichment of common macrophage markers such as *Cd68*, and additional genes typically associated with immunosuppressive TAMs or M2-like macrophages including *Arg1*, *Hmox1*, *Msr1*, *Trem2*, *Spp1* and *Ctsk* (**Fig. 5D, Supplementary** Fig. 5E) ^59–67^. To validate this, we performed IHC and observed higher levels of MSR1 (CD204), HMOX1 and ARG1 expression in LUSCs compared to LUADs (**Fig. 5E**). In contrast, the oncogenic driver in this mouse model, SOX2, was expressed in both LUSCs and LUADs. These findings demonstrate that *Tmprss11b* expression correlates with infiltration of immunosuppressive M2-like macrophages.

**Figure 5.**
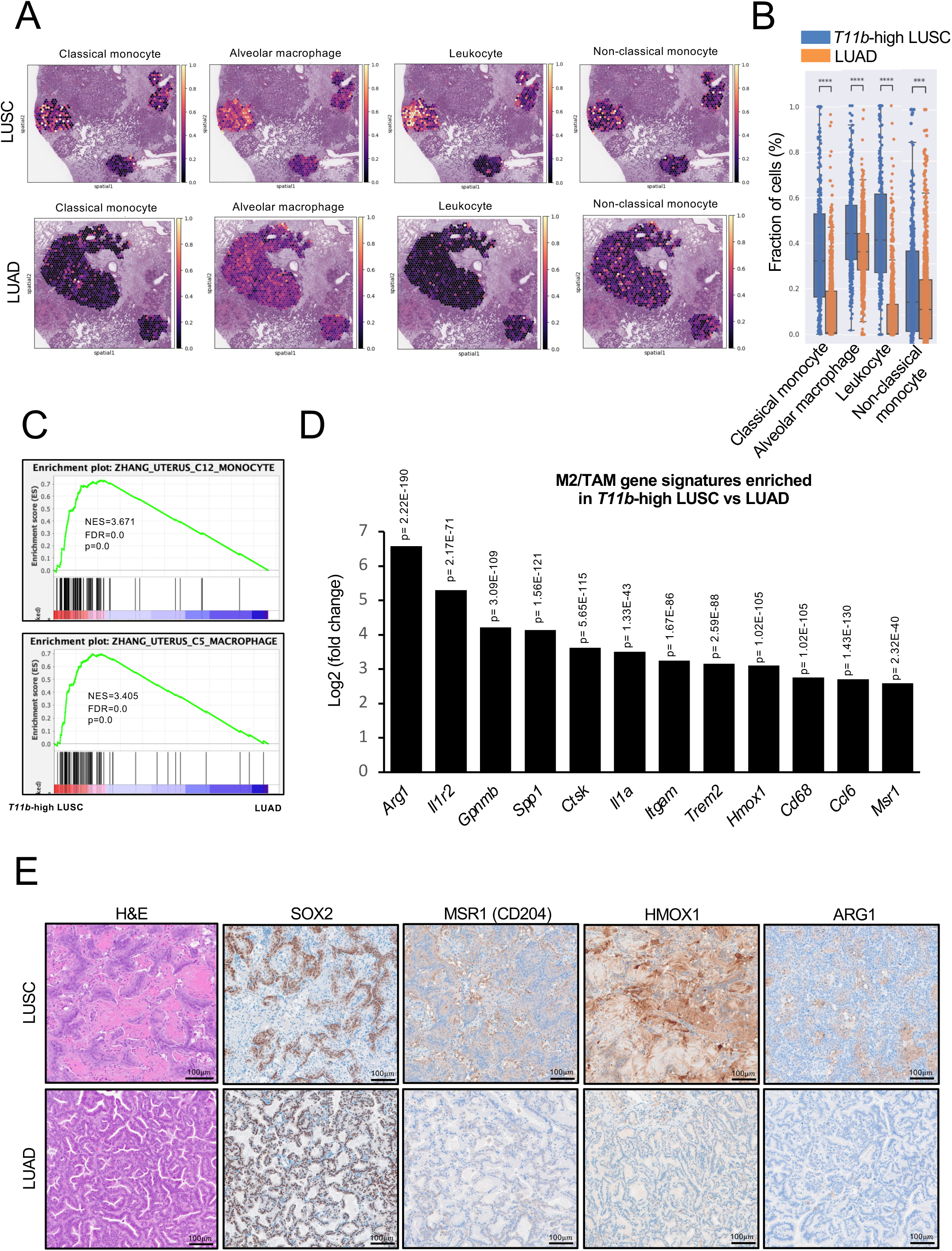
Infiltration of M2-like macrophages in *Tmprss11b*-high squamous tumors. **A)** Spatial plots from cell deconvolution analysis showing the distribution of various immune cell populations in LUSCs (top) and mucinous LUADs (bottom). **B)** Quantification of immune cell populations in (A). **C)** Gene set enrichment analysis (GSEA) of *Tmprss11b*-high LUSC versus LUAD spatial transcriptomics data with normalized enrichment scores (NES), false discovery rate (FDR) and p values for the indicated gene signatures. **D)** Bar graph representing the top M2-like/TAM genes from the differential gene expression (DEG) analysis of *Tmprss11b*-high LUSC versus LUAD spatial transcriptomics data. **E)** Representative H&E images and immunohistochemistry (IHC) for SOX2, MSR1 (CD204), HMOX1 and ARG1 in SNL LUSC (top) and mucinous LUAD (bottom).

### *Tmprss11b*-high squamous tumors and acidified regions of the TME are enriched for immunosuppressive macrophages

Given the role of TMPRSS11B in promoting lactate export and the ability of lactate to promote polarization of macrophages to the M2-subtype ^30–33^, we hypothesized that the infiltration of M2-like macrophages in *Tmprss11b*-high squamous tumors was associated with elevated levels of lactate in the tumor microenvironment. To investigate this, we utilized ultra pH-sensitive (UPS) nanoparticles conjugated to the indocyanine green (ICG) fluorophore (**Supplementary** Fig. 6A) ^68^. We utilized a UPS nanoparticle with a threshold pH of 5.3 that disassembles and electrostatically binds to the surrounding tissue when the pH is below 5.3 ^69^. The illuminated ICG signal defines the regions of increased acidity that are driven by secretion of lactic acid. We administered the UPS nanoparticles to control animals (without Cre) and tumor bearing SNL mice (with Cre) by tail vein injection and performed *ex-vivo* imaging of the lungs (**Supplementary** Fig. 6B). We observed high ICG signal in the lungs of tumor-bearing SNL mice administered with Ad-Cre, indicating increased acidification (**Fig. 6A-B**). Moreover, the ICG signal was adjacent to regions expressing *Tmprss11b* and macrophage markers, including *Cd68* and *Spp1* (**Fig. 6C**). Consistent with this, our prior data showed the highest level of acidity at the tumor and stromal interface, where cancer cells secrete lactic acid into the stromal areas, leading to the nanoprobe activation and internalization by stromal cells ^69^. We performed GSEA and observed an enrichment of pathways associated with macrophages and monocytes, and oncogenic signaling in regions with low pH (**Fig. 6D, Supplementary** Fig. 6C). Cell deconvolution analysis further revealed an enrichment for select immune cell subtypes, including monocytes and macrophages, in low pH regions (**Supplementary** Fig. 6D). Finally, DEG analysis and IHC validation revealed the expression of M2-like and TAM gene signatures (**Fig. 6E**), and expression of HMOX1 and MSR1 in regions with high UPS signal, respectively (**Fig. 6F**). Taken together, these findings reveal the presence of acidified regions in lung squamous tumors that accumulate immunosuppressive M2-like macrophages.

**Figure 6.**
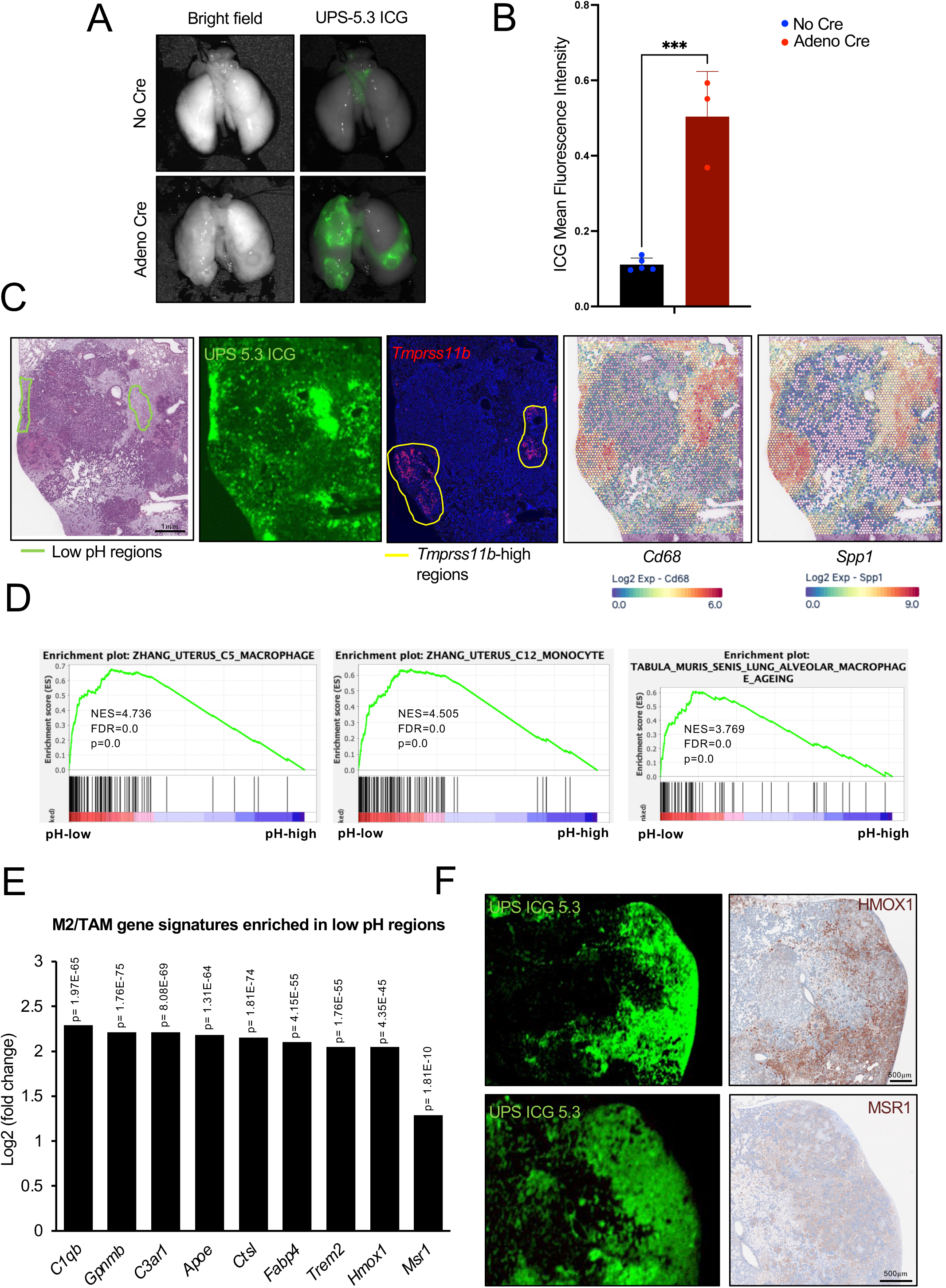
*Tmprss11b*-high squamous tumors and acidified regions of the TME are enriched for immunosuppressive macrophages. **A)** Representative *ex-vivo* images of lungs from SNL mice, control (no Cre) or infected with Ad-Cre (11 months post infection), 24hrs post injection with PDBA ICG 5.3 ultra pH sensitive (UPS) nanoparticle, along with the quantification of the ICG signal (control n=5 mice; Ad-Cre n=3 mice). Bright field images (left) and the corresponding ICG fluorescence images (right) are shown. **B)** Quantification of mean ICG fluorescence intensity (n=3 mice per group). **C)** H&E image of a SNL lung section used for spatial transcriptomics, with ICG signal on the same section, and RNAscope of *Tmprss11b* (red) on a serial section. Also shown are spatial plots depicting enrichment of *Cd68* (macrophage marker) and *Spp1* (immunosuppressive macrophage marker) transcripts in these regions. **D)** Gene set enrichment analysis (GSEA) of immune cell pathways from the low pH versus high pH spatial transcriptomics data with normalized enrichment scores (NES), false discovery rate (FDR) and p values for the indicated gene signatures. **E)** Bar graph representing top M2-like/TAM genes from the differential gene expression (DEG) analysis of low pH versus high pH spatial transcriptomics data. **F)** Representative ICG fluorescence images (left) and the corresponding immunohistochemistry (IHC) validation of HMOX1 and MSR1 (right) in SNL lung sections.

### *Tmprss11b*-high squamous tumors and low pH regions have elevated levels of lactate

Given the observed enrichment of immunosuppressive M2-like macrophages in *Tmprss11b*-high squamous tumors and the surrounding TME, we reasoned that higher levels of lactate should be present in the acidified regions. To demonstrate that regions with low pH and increased acidification in *Tmprss11b*-expressing tumors accumulate lactate, we performed laser capture microdissection (LMD) and mass spectrometry to quantify metabolites in the lungs from SNL mice (**Fig. 7A**). We used H&E staining to annotate regions of LUSC, LUAD, and normal lung. The ICG signal was used to annotate regions of low pH (**Fig. 7B, Supplementary** Fig. 7A). Quantification of metabolite levels revealed a significant enrichment of lactate in lung squamous cell carcinomas (LUSC) compared to lung adenocarcinomas (LUAD) and surrounding normal lung tissues (**Fig. 7C**). Moreover, we observed elevated lactate levels in low pH regions identified with the ultra pH sensitive nanoprobe (**Fig. 7D**). Collectively, these data demonstrate that LUSC tumors acidify the TME due to their high expression of *Tmprss11b*, resulting in an immunosuppressive environment that enhances tumor growth.

**Figure 7.**
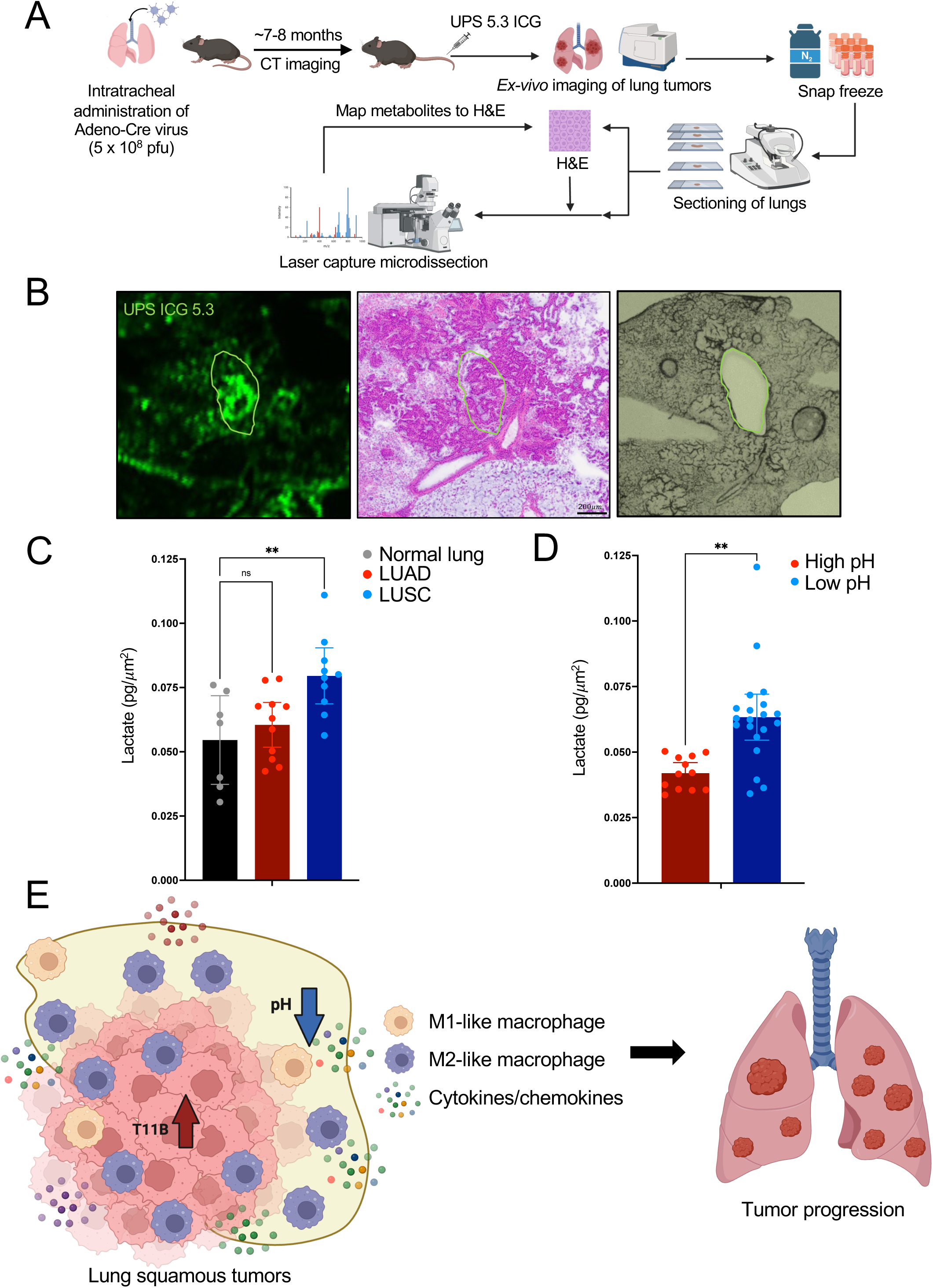
*Tmprss11b*-high squamous tumors and acidified regions have elevated levels of lactate. **A)** Schematic representation of the experimental pipeline to assess metabolites in SNL lung tumors through laser capture micro-dissection (LMD) followed by mass spectrometry. **B)** Representative ICG fluorescence images (depicting nanoparticle accumulation) (green), H&E image (middle), and bright field image of the serial section (right) used for LMD with the same region annotated in green. **C)** Quantification of lactate concentration (pg/μm^2^) in SNL lung tissues; normal, LUADs and LUSCs (n=7-11 regions per group from a total of 2 mice). Ordinary one-way ANOVA with Dunnett’s multiple comparisons test was used for the statistical analysis. **D)** Quantification of lactate concentration (pg/μm^2^) in the regions of low acidity (high pH) and high acidity (low pH), in the SNL lung tissues (n=12-20 regions per group from a total of 2 mice). Ordinary one-way ANOVA with Dunnett’s multiple comparisons test was used for the statistical analysis. **E)** Schematic representation of the immunosuppressive niche established in *Tmprss11b*-high lung squamous tumors and the surrounding acidified regions in the tumor microenvironment.

## Discussion

Based on our prior studies demonstrating a role for TMPRSS11B in promoting tumorigenesis and enhancing lactate export *in vitro*, we hypothesized that TMPRSS11B expression would promote an immunosuppressive tumor microenvironment *in vivo*. Indeed, loss of TMPRSS11B in a syngeneic mouse model confirmed the importance of TMPRSS11B in LUSC tumorigenesis and identified alterations in immune cell populations and signaling upon depletion of *Tmprss11b*. Furthermore, using the *Rosa26-Sox2-Ires-Gfp^LSL/LSL^*;*Nkx2-1^fl/fl^*;*Lkb1^fl/fl^*(SNL) GEMM model of LUSC, we observed specific expression of TMPRSS11B in lung squamous tumors. A similar gene expression pattern is observed in the LP (*Lkb1*^fl/fl^, *Pten^f^*^l/fl^) GEMM model, where *Tmprss11b* is one of the top upregulated genes in lung squamous tumors ^70^. Leveraging the SNL model, we performed spatial transcriptomics on lung tumors, which revealed increased oncogenic signaling and accumulation of immunosuppressive M2-like macrophages in *Tmprss11b* expressing squamous tumors. We assessed the *in vivo* acidification of squamous tumors using ultra pH sensitive nanoparticles and uncovered the presence of low pH regions adjacent to squamous tumors that are rich in immunosuppressive M2-like macrophages. Given the ability of lactate to promote macrophage polarization and immunosuppression ^30–33^, we hypothesized that lactate is present in these regions. Accordingly, laser capture microdissection followed by mass spectrometry demonstrated that regions with low pH exhibited elevated lactate abundance. Taken together, these results support a model whereby TMPRSS11B expression in LUSC results in acidification of the TME due to enhanced lactate export. This produces an immunosuppressive environment, characterized by the infiltration of M2-like macrophages, which promotes tumor progression (**Fig. 7E**).

Treatment modalities for LUSC patients have significantly advanced in recent years. Immunotherapy is approved as a first-line therapy and has shown promising improvement in progression free survival, either as a monotherapy or in combination with chemotherapy. However, only a subset of the patients respond with durable benefit and patients often develop toxicity or resistance to immunotherapy, warranting the need for alternative therapies ^6,71,72^. Additionally, efforts to target frequently upregulated molecules in LUSC such as FGFR, PI3K and CDK4/6 have been largely unsuccessful ^6^. Collectively, these data suggest the need for identification and development of new targeted therapies for molecules specifically upregulated in LUSC. The selective upregulation of TMPRSS11B in lung squamous tumors and the impact on immune suppression and oncogenic signaling pathways nominates this cell surface enzyme as a promising therapeutic target.

Our studies suggest that the enrichment of TMPRSS11B in lung squamous tumors may be attributed to its expression in hillock cells. The unique properties of this cell type include their ability to form stratified structures with a gradient of KRT13 and TP63 expression. This results in upper layers of ciliated luminal (termed hillock luminal) cells marked by expression of KRT13 and underlying layers of basal stem (termed hillock basal) cells marked by high expression of TP63^16,17,48^. The hillock basal cells appear to be more proliferative than the basal cells of the pseudostratified epithelium while the hillock luminal cells consist of tightly interlocked squamous cells that form a protective barrier for the underlying stem cells. Hillocks cells are highly resistant to insults including chemical injury and viral infection ^16^. Furthermore, hillock cells express genes involved in cellular adhesion and immunomodulation, including *Cldn3*, *S100a11*, *Ecm1*, *Lgals3* and *Anxa1* ^17,48–53^. Interestingly, recent studies have suggested an important role for members of the type-II transmembrane serine protease (TTSP) family in barrier function ^73^. Given the unique enrichment of TMPRSS11B in hillock-like cells, it is plausible that TMPRSS11B contributes to barrier function and immune regulation mediated by hillock cells.

A recent study suggested hillock cells as precursors of LUSC ^16^. Our companion study suggests that KLF4 can induce hillock gene signatures in human bronchial epithelial cells (Izzo et al., 2025, co-submitted). Taken together with our data demonstrating that KLF4 induces *Tmprss11b*, this suggests a model whereby KLF4 is transiently upregulated in hillock basal cells, giving rise to suprabasal hillock luminal cells that express KRT13 and TMPRSS11B. This population of cells is hypothesized to function as a barrier, protecting the underlying hillock basal stem cells from various insults. The potential dysregulation of signaling in hillock cells that results in the progression to squamous tumors and the role of TMPRSS11B in this process is not completely understood. Future studies are warranted to fully elucidate the role of TMPRSS11B in the hillock cell state and identify substrates of this serine protease in normal lung development and in LUSC.

Macrophages comprise a significant fraction of tumor associated immune cells in the lung and have been shown to influence tumor growth and treatment outcomes ^74–76^. Additionally, the presence of the classically activated M1 subtype, which are anti-tumorigenic and the alternatively activated M2 subtype, which are immunosuppressive and tumor promoting, adds an additional layer of complexity to understanding the impact of macrophages in the TME. Higher levels of M2-like macrophages are associated with a decrease in overall survival (OS) of LUSC patients ^77,78^. Our studies have uncovered an enrichment of M2-like macrophages in *Tmprss11b*-high squamous tumors and the surrounding acidified regions. In addition, we identified higher lactate levels in these regions. Taken together with our previously identified role for TMPRSS11B in promoting lactate export and the reported role of lactate in polarization of macrophages to the M2 subtype, these results suggest an association of TMPRSS11B with the recruitment or production of immunosuppressive M2-like macrophages. Recent findings have also implicated a role for IL6-JAK-STAT3 signaling in promoting polarization of macrophages to the M2 subtype ^79^. Interestingly, our studies uncovered upregulation of an ‘IL6_JAK_STAT3_Signaling’ pathway signature associated with increased *Tmprss11b* expression, further hinting at a potential role for TMPRSS11B in promoting macrophage polarization. Additional studies are needed to fully elucidate the link between TMPRSS11B and macrophage polarization and to determine whether this is solely due to the impact on lactate metabolism or whether additional functions of this enzyme play a role.

Collectively, these findings shed light on an important metabolic regulator of lung squamous cell carcinoma. Given our demonstration that TMPRSS11B promotes tumor growth and an immunosuppressive TME, the development of monoclonal antibodies or small molecule inhibitors that target this enzyme at the surface of LUSCs represents an exciting avenue for development of new therapeutics for this deadly malignancy.

## Materials and Methods

### Mice

*Rosa26-Sox2-Ires-Gfp^LSL/LSL^*;*Nkx2-1^fl/fl^*;*Lkb1^fl/fl^*(SNL) mice were provided by Dr. Trudy Oliver (Duke University). The mice were maintained on a mixed background through intercrosses. DBA/2 mice were purchased from The Charles River Laboratory.

### Ethics Statement

Mice were monitored closely throughout all experimental protocols to minimize discomfort, distress or pain. If any signs of pain and distress were detected (disheveled fur, decreased feeding, significant weight loss (>20% body mass), limited movement or abnormal gait), the animal was removed from the study and euthanized. All procedures involving mice were performed in accordance with the recommendations of the Panel on Euthanasia of the American Veterinary Medical Association and protocols approved by the UTSW Institutional Animal Care and Use Committee.

### Cell culture

KLN205 murine lung squamous cell carcinoma cells (from Dr. Rolf Brekken) were cultured in Gibco^TM^ DMEM media (Cat: 11995065) supplemented with 10% FBS and 1% antibiotic and antimycotic (Cat: 15240062).

### Plasmids

LentiCRISPR version 2, PAX2 and MD2 plasmids were obtained from Addgene (52961, 12260 and 12259). sgRNA sequences targeting *Tmprss11b* were selected from the Brie sgRNA library (target gene ID: 319875). The SMARTvector inducible lentiviral shRNA plasmids were obtained from Dharmacon. The sgRNAs and shRNAs sequences are provided in **Supplementary Table 2**.

### Generation of *Tmprss11b* knockout cells

HEK293T (1 x 10^8^) cells were co-transfected with lentiCRIPSR version 2 (1 μg), PAX2 (666 ng) and MD2 (333 ng) helper plasmids using Effectene Transfection Reagent (Qiagen, Cat: 301425). Lentiviral supernatant was collected 48 hours post-transfection and filtered. Recipient KLN205 cells were infected with viral supernatant containing 8 μg/ml polybrene (Sigma-Aldrich) and replenished with fresh media. After 48 hours, the transduced cells were cultured in fresh media containing 2 μg/ml puromycin for 7-9 days.

### Generation of inducible *Tmprss11b* knockdown cells

HEK293T (1 x 10^8^) cells were co-transfected with SMARTvector inducible lentiviral shRNA plasmid (1 μg), PAX2 (666 ng) and MD2 (333 ng) helper plasmids using Effectene Transfection Reagent (Qiagen, Cat: 301425). Lentiviral supernatant was collected 48 hours post-transfection and filtered. Recipient KLN205 cells were infected with viral supernatant containing 8 μg/ml polybrene (Sigma-Aldrich) and replenished with fresh media. After 48 hours, transduced cells were selected in fresh media containing 2 μg/ml puromycin for 7-9 days and cultured in 2-3 μg/ml doxycycline. shRNA expression was monitored over 4 days using the turbo RFP expression. At 4 days post-induction, the cells were harvested for RNA to assess the knockdown.

### Tumorigenesis assays

KLN205 cells (3 x 10^5^) expressing non-targeting sgRNA or *Tmprss11b* sgRNA were injected subcutaneously into the right flanks of 6-to 8-week-old DBA/2 female mice (Charles River). Tumor volumes were measured every 3 days using calipers until the average tumor mass reached 600 mm^3^. sgRNA sequences are provided in **Supplementary Table 2**.

For inducible knockdown, KLN205 cells (5 x 10^5^) expressing scrambled shRNA or *Tmprss11b* shRNA were injected subcutaneously into the right flanks of 6- to 8-week-old DNA/2 female mice (Charles River). Tumor volumes were measured every 3 days using calipers. When the average tumor volume across all the study groups reached 100 mm^3^, mice were maintained on doxycycline water (2g/L doxycycline and 2% sucrose) for the duration of the experiment.

Tumor volumes were calculated using the formula (length x width^2^)/2. At the terminal timepoint, the tumors were resected for downstream analysis.

### RNA extraction and quantitative real-time PCR (qRT-PCR) analysis

Total RNA was isolated from cells using RNeasy Mini Kit (Qiagen). Total RNA was isolated from tumor tissues using Trizol (Ambion by Life Technologies; 15596018), followed by additional cleanup and DNase digestion using RNeasy Mini Kit (Qiagen). For qRT-PCR of mRNA, complementary DNA (cDNA) synthesis was performed with 1-5 μg of the total RNA for reverse transcription using Superscript IV VILO Master Mix (5x) (Invitrogen, Cat: 11756050). mRNA expression was assessed using TaqMan probes (Invitrogen) corresponding to *Tmprss11b* (Mm00621706_m1, Mm00621702_m1 and Mm00621704_m1) and *Gapdh* (Mm99999915_g1) and calculated using the 2^ddCt^ method.

### RNA-Sequencing and analysis

RNA was isolated from tumor tissues as indicated in the last section. The sequencing was performed at the McDermott Center Next Generation Sequencing Core at UT Southwestern Medical Center and the analysis was performed at the McDermott Center Bioinformatics lab. Fastq files were quality checked using fastqc (v0.11.2)^80^. Read from each sample were mapped to the reference genome using STAR(v2.5.3a)^81^. Read counts were generated using featureCounts ^82^ and the differential expression analysis was performed using edgeR ^83^.

### Intratracheal administration of adenovirus and tissue collection

Cre-expressing adenovirus (Ad-Cre) was purchased from Viral Vector Core Facility (University of Iowa). *Rosa26-Sox2-Ires-Gfp^LSL/LSL^*;*Nkx2-1^fl/fl^*;*Lkb1^fl/fl^*(SNL) mice (males and females) 6- to 8-weeks of age were intratracheally administered Ad-Cre at 1-10 x 10^8^ pfu/mouse.

Mice were euthanized by intraperitoneal administration of an overdose of Avertin at the indicated time-points. Lungs were inflated and perfused through the trachea with 4% paraformaldehyde (PFA), fixed overnight, washed in 1X PBS for 24-48h, then transferred to 70% ethanol and subsequently embedded in paraffin. Sections were cut and stained with H&E by the UTSW HistoPathology Core.

### Immunohistochemistry

The formalin fixed paraffin embedded (FFPE) slides were de-paraffinized with xylenes and hydrated with ethanol washes. Slides were then treated with either Citrate-based (pH 6.0) or Tris-based (pH 9.0) Antigen Unmasking Solution (Vector Lab, H-3300 or H3301). Antigen retrieval was performed using a steamer, followed by blocking using BLOXALL^TM^ Endogenous Blocking Solution (Vector Lab, SP-6000) and washed in 1X PBS. Then the slides were incubated with 2.5% Normal Goat Serum (Vector Lab, S1012) followed by incubation with appropriate primary antibodies diluted in 2% bovine serum albumin (BSA) (in PBS) at 4°C overnight. After extensive washing with either PBS/TBS, the slides were incubated with ImmPRESS ® HRP Goat Anti-Rabbit IgG (Vector Lab, MP7451) or Goat Anti-Rat IgG (Vector Lab, MP7404) at room temperature (RT) for 30 minutes. The signal was developed with ImmPACT ® DAB Substrate (Vector Lab, SK-4105), and sections were counterstained with hematoxylin (Vector Lab, H-3404), and mounted using Prolong^TM^ Diamond Antifade Mountant (Invitrogen, Cat: P36965). Whole slide images were captured using Hamamatsu NanoZoomer S60 at the UTSW Whole Brain Microscopy Facility. Antibody information is provided in **Supplementary Table 1**.

### Multiplex fluorescent Immunohistochemistry and quantification

The fluorescent immunohistochemistry was performed using Akoya Opal 3-Plex Detection kit (NEL810001KT) following manufacturer’s instructions. Briefly, FFPE slides were de-paraffinized using xylenes, hydrated with ethanol washes and then fixed with 4% paraformaldehyde (PFA). Antigen retrieval was performed at either pH 6.0 or pH 9.0 using a microwave. Blocking was performed using Antibody Diluent/Block from the kit. Slides were then incubated with the primary antibodies at 4°C overnight. After extensive washing with TBST, the slides were incubated with Opal Anti-Ms+Rb HRP and then the appropriate Opal/TSA plus fluorophore. For multiplexing, this was followed by another round of antigen retrieval and same steps were followed as before until all the targets have been achieved. The slides were then incubated with Spectral DAPI, followed by mounting with Prolong^TM^ Diamond Antifade Mountant (Invitrogen, Cat: P36965). Whole slide images were captured using Zeiss Axioscan.Z1 at the UTSW Whole Brain Microscopy Facility.

For quantification, the Zen Blue software from Zeiss was used. For each slide, 10-14 equal sized fields are selected across the slide and mean fluorescent intensities (MFI) for each channel was calculated for each field.

### RNA in-situ hybridization

RNA in-situ hybridization was performed on formalin-fixed paraffin-embedded (FFPE) tissue sections following instructions from the RNAscope Multiplex Fluorescent Kit v2 (Cat. No. 3231100, Advanced Cell Diagnostics) using the following probes purchased from Advanced Cell Diagnostics: *Tmprss11b* (Cat No. 1268741-C2), *Sox2* (Cat No. 401041-C3) and negative control *DapB* (Cat No. 310043-C2). Slides were counterstained with DAPI (Cat No. 320858, Advanced Cell Diagnostics) and mounted with Prolong^TM^ Diamond Antifade Mountant (Invitrogen, Cat: P36965). Whole slide images were obtained using Axioscan.Z1 and Vectra Polaris^TM^ imagers.

### Ultra pH-sensitive (UPS) nanoparticle imaging

Ultra pH-sensitive (UPS) nanoparticles were used for imaging acidic regions in mouse lung tumors and other tissues ^84^. Briefly, SNL mice bearing lung tumors (from CT imaging) were injected with 2.5mg/kg of UPS nanoparticle PDBA-ICG 5.3. 24 hours post-infection, the mice were euthanized by intraperitoneal administration of an overdose of Avertin. Lungs were inflated and perfused through the trachea with 4% paraformaldehyde (PFA). The lungs were then imaged with liver, spleen, kidney, heart and brain from the same mouse for brightfield (BF) and Indocyanine green (ICG) signals using Pearl ® Trilogy small animal imaging system (LICOR Bio). Post imaging, the lungs were fixed overnight in 4% PFA, washed in 1X PBS for 24-48h, then transferred to 70% ethanol and subsequently embedded in paraffin. Sections were cut and stained with H&E. The ICG signal in lungs were quantified as mean fluorescent intensity (MFI) using the Image Studio^TM^ software (LICOR Bio). For the analysis, ICG signal in the brain was used as background and normalized for lungs and other tissues accordingly.

### MRI and CT imaging

Magnetic resonance imaging (MRI) was performed on mice using the 7T Bruker Biospec instrument at the Advanced Imaging Research Center at UT Southwestern with the help of Dr. Janaka Wansapura.

Computed tomography (CT) imaging was performed on mice using the X-Cube CT instrument from Molecubes at the Preclinical Radiation Core Facility at UT Southwestern.

### Surveyor Assay

Surveyor assay was used to assess the efficiency of the different *Tmprss11b* targeting sgRNAs. The assay was performed on *Tmprss11b* knockout and non-targeting control KLN205 cells using Guide-it^TM^ Mutation Detection kit (Takara, Cat No. 631448) following manufacturer’s protocol. Briefly, genomic DNA was isolated from the above-mentioned cells, and the mutated regions were amplified using specifically designed surveyor primers (provided in **Supplementary Table 2**). Then the amplified sequences were subjected to cleavage by the Guide-it resolvase and screened by running samples on agarose gel electrophoresis.

### Spatial transcriptomics: Visium CytAssist

Hematoxylin and eosin-stained tissue slides were scanned at 40x magnification using the Leica Biosystems Aperio AT2 DX brightfield slide scanner. Regions of lung adenocarcinoma and lung squamous cell carcinoma morphology were annotated. The regions of interest for spatial transcriptomics were digitally annotated using QuPath (v.5.0.1) and exported as a TIFF image file.

Tissue slides were decoverslipped, washed, and rehydrated following the 10x Genomics Visium CyAssist Tissue Preparation Guide (CG000518 rev. B). Slides were immersed in xylene then placed on a metal block cooled with dry and the coverslip was removed with a razor blade. Sections were washed and rehydrated by sequential immersions in xylene, 100% ethanol, 96% ethanol, and 70% ethanol, according to manufacturer recommendations. A region of the tissue outside of the region of interest was cut away for RNA quality control. RNA was isolated with the Qiagen RNeasy Mini Kit (cat: 74104), and quality was evaluated using the Agilent Technologies RNA 6000 Pico kit (cat: 5067-1513) with a DV200 assay in the Agilent 2100 BioAnalyzer.

Rehydrated samples were destained and decrosslinked following 10x Genomics protocol CG000520 rev B. Sample preparation was performed with a 10x Genomics Visium CytAssist Spatial Gene Expression for FFPE, Mouse Transcriptome, 6.5mm kit (PN-1000521) and its corresponding protocol (GC000495 rev. C). Mouse whole-transcriptome left and right-hand probes were hybridized to the tissue at 50°C for 18 hours. Hybridized probes were ligated at 37°C for 60 min. Tissue was then stained with 10% eosin Y for one minute and rinsed with 1X PBS. Slides were placed in the Visium CytAssist instrument and manually aligned for capture of the regions of interest. Gene expression probes were released from tissue and captured by the Visium CytAssist spatial slide using the CytAssist instrument at 37°C for 30 min. The spatial slide was washed with 2X SSC, and captured probes were extended on the slide at 45°C for 15 min. Probe amplicons were eluted from the capture slide with 0.08M KOH and neutralized with Tris-HCl pH 8.0. Samples probes were pre-amplified by PCR, and optimal cycle determination was performed using qPCR. Final sequencing libraries were prepared using 10x Genomics dual index plate TS set A (PN-3000511) and sequenced with read depth of at least 50,000 reads per spot using the Singular Genomics G4 platform with an F3 flow cell.

### Spatial transcriptomics analysis pipeline

Visium spatial transcriptomics reads were processed using 10x Genomics Space Ranger v2.0.1. Samples were post-processed using R v.4.3.1 ^85^ with Seurat v. 5.0.3 ^86^. Spots corresponding to Adenocarcinoma, Squamous cell carcinoma, and low pH were annotated using Loupe Browser v.6.5.0 based on morphology from H&E images and corresponding assays in adjacent tissue sections. Samples were processed for quality control, preserving spots with a minimum of 700 read counts and 200 genes, and genes present in at least 5 spots. RNA gene expression libraries were normalized using SCtransform ^87^.

Spots expressing *Tmprss11b* at or above 95th percentile were assigned “Tmprss11b-High”. Squamous tissue regions were further subclassified according to dichotomized *Tmprss11b* expression. All spots were categorized into “*Tmprss11b*-High Squamous”, “Adenocarcinoma”, “Non-Tumor Lactate-High”, “Non-Tumor Lactate-Low”, and “Other” classes. Differential gene expression among all these annotated regions was calculated using Seurat’s FindAllMarkers function, and direct comparisons between two classes was performed with Seurat’s FindMarkers function. Differentially expressed genes were used to perform pathway enrichment analysis using clusterProfiler v.4.10.0 ^88,89^ using Gene Ontology^90,91^, KEGG references ^92,93^. Gene expression signatures for “Mouse_Hillock”, “Mouse_Basal”, and “Mouse_Mucinous phenotypes were evaluated for each spot using Seurat’s AddModuleScore function.

### Tangram cell type deconvolution

Spatial transcriptomics technologies provide spatially resolved gene expression data; however, their resolution is often lower than that of single cell RNA sequencing (scRNA-seq). Consequently, each spatial spot or voxel may contain contributions from multiple cell types, making it challenging to determine the precise cellular composition of a tissue. Spatial deconvolution addresses this issue by integrating high-resolution scRNA-seq data with lower-resolution spatial transcriptomics data to infer the relative proportions of different cell types at each spatial location. Among the available spatial deconvolution methods ^94–97^ , we employed Tangram ^94^ due to its ability to map single cells onto spatial transcriptomics data across multiple platforms with high accuracy. Tangram utilizes deep learning-based optimal transport to achieve unbiased alignment, multimodal integration, and gene-level precision. By leveraging both scRNA-seq and spatial transcriptomics datasets, Tangram optimally assigns single cells to spatial locations while preserving gene expression consistency across modalities.

Data preprocessing: We obtained reference single-cell RNA sequencing (scRNA-seq) data from The Tabla Muris Consortium (*Nature* 2018) ^98^ and spatial transcriptomics (ST) data from relevant datasets. The scRNA-seq dataset was preprocessed by normalizing gene expression counts and performing dimensionality reduction using principal component analysis (PCA). Cells were then clustered based on their transcriptional profiles, and each cluster was annotated with its corresponding cell type.

Mapping scRNA-seq to spatial transcriptomics: Tangram applies a probabilistic framework to align single-cell gene expression profiles with spatial transcriptomics data by optimizing the mapping based on a reconstruction loss function. The spatial mapping was performed using the following steps:

1. Model Initialization: The single-cell and spatial gene expression matrices, along with cell-type annotations, were input into Tangram, ensuring that only genes common to both datasets were retained.
2. Optimization: Tangram estimated the optimal mapping by solving a constrained optimization problem that minimizes the reconstruction error while preserving gene expression consistency between single-cell and spatial transcriptomics data.
3. Cell-Type Decomposition: Using the trained model, Tangram inferred the proportion of each cell type at individual spatial locations, generating a probabilistic estimate of cellular distributions across the tissue.

### Laser Capture Microdissection

The SNL mice administered with Ad-Cre were euthanized at 8-months post infection by intraperitoneal administration of an overdose of Avertin. The lungs were flash frozen and sectioned into ∼ 50 µm thick sections on polyethylene terephthalate membrane (PTFE) slides. Leica laser LDM6 microdissection microscope was used to selectively cut the tissue areas exclusively identified as lung squamous cell carcinoma, lung adenocarcinoma, adeno-squamous, low pH, high pH and normal tissue regions based on adjacent histology staining and fluorescent UPS probe distribution. Each excised area was collected in individual PCR vials. 100µL of 0.1% formic acid in water was added to each PCR vial to extract water soluble metabolites. The vials were then subjected to sonication for 30 min in a water bath followed by 20 min vortex at 2,500 rpm (VWR DVX-2500 multi-tube vortex mixer). The samples were centrifuged at 14000 xg for 10min. Then 45µL of the supernatant was transferred into an autosampler vial to mix with 5µL internal standard (IS) mix (5 µg/mL). The sample injection volume was 10µL.

A Shimadzu CBM-20A Nexera X2 series LC system (Shimadzu Corporation, Kyoto, Japan) equipped with degasser (DGU-20A) and binary pump (LC-30AD) along with auto-sampler (SIL-30AC) and (CTO-30A) column oven was used. Chromatographic separation of glucose, lactic acid, pyruvic acid, glutamic acid, succinic acid and citric acid were achieved using a Phenomenex Luna C8 (2), 5µm 100 Å, 150×2.0 mm column. 0.1% formic acid in water was used for mobile phase A, and 0.1% formic acid in acetonitrile was used as mobile phase B. The LC flow rate was set at 0.3 mL/min with gradient started from 3% of B for 1 min, 30% B by 5 min, 98% B by 5.5 min maintained to 7 min, then switched back to 3% B by 7.1 min and maintained to 8 min. The autosampler was maintained at 5 °C. Injection volume of 10 µL was used.

The primary stock solutions for all analytes including standards and isotope labeled internal standards (IS) (1.0 mg/mL) were prepared in LC/MS grade water and subsequent dilutions were prepared in 0.1% formic acid in water. Standard curves were prepared at a range of concentrations at 1, 5, 10, 25, 50, 100, 500, 1000, 5000 and 10000 ng/mL with different lowest calibration concentration points suited for different metabolites. IS mix of 5ug/mL of each D-glucose (U-^13^C_6)_, L-Lactic acid-^13^C_3,_ Pyruvic acid sodium salt ^13^C_1_, L-Glutamic acid-^13^C_5_, Succinic acid −2,2,3,3-D4, and Citric acid 2,2,4,4-D4 was also prepared in 0.1% formic acid in water. 5µL of IS mix (5 µg/mL) was spiked in 45 µL of standard curves and samples to correct for any response-based differences created from the instrument or sample preparation.

An AB Sciex (Foster City, CA, USA) 6500+ QTRAP mass spectrometer, equipped with a Turbo ion spray™ (ionization source) was used as the detector. The mass spec interface temperature was set at 500°C. The ion spray voltage was set at −4500 Volts. Other parameters such as nebulizer gas, curtain gas, auxiliary gas and CAD gas were set at 50, 55, 65 and Medium, respectively. Detection of the ions was performed in multiple reaction monitoring (MRM) mode, the transition pairs Q1/Q3 were set on unit resolution. MRM transition pairs and each of their entrance potential (EP), collision energy (CE), and collision exit potential (CXP) are presented in Supplementary table 4.

The LC/MS/MS data were processed by Analyst software (version 1.7.3). The results were fitted to linear regression analysis using 1/*X*^2^ as weighting factor. Quantifier ions Q3 are listed in **Supplementary Table 4.** The final concentration of metabolites was calculated by normalizing the measured amounts to the area of the dissected tissue regions.

### Functional analysis of gene sets

Pathway and network analysis were performed using GSEA 4.3.2 application from the Broad Institute ^99,100^. The GSEA Preranked tool was used for ranked gene list using the rank_score=sign_of_FC*-log(pval) for all the expressed genes in the RNA-seq dataset with a weighted scoring scheme and using rank_score=log2FC for all the genes showing significant differences in expression (pval <0.05) in the spatial transcriptomics dataset with a weighted scoring scheme.

### Statistics and reproducibility

An unpaired *t*-test with Welch’s correction was used for comparisons between two groups (for comparing MFI from IHC-F for CD4+ T cells and other indicated analyses). Ordinary one-way ANOVA with Dunnett’s multiple comparisons test or Brown-Forsythe and Welch ANOVA test with Dunnett’s T3 multiple comparisons test was used for comparisons between more than two groups (for qRT PCR analyses, comparing tumor volumes from implantation studies and other indicated analyses). For the tumorigenesis assays in syngeneic mice, linear mixed-effects models were used to investigate if there were significant differences in tumor volume over time among the three groups. A Wilcoxon Rank Sum test was used to compare the distribution of *Tmprss11b* expression across tissue regions.

## Data Availability

Spatial transcriptomics data that support the findings of this study have been deposited in the Gene Expression Omnibus (GEO) under accession code GSE292706. The accession number for the RNA sequencing data is GSE292085.

## Acknowledgements

We thank Joshua Mendell and members of the O’Donnell laboratory for critical reading of the manuscript. K.A.O. is supported by the NCI (R01CA273585, R01CA207763, and P50CA70907), the Cancer Prevention Research Institute of Texas (CPRIT RP190610, RP200327, and RP250391), the Welch Foundation (I-1881), the V Foundation (T2021-011), and the Department of Defense (DoD LC190249). We also thank the UTSW Tissue Management Shared Resource, a shared resource at the Simmons Comprehensive Cancer Center, which is supported in part by the National Cancer Institute P30 CA142543. Slide scanning was made possible on Zeiss Axioscan.Z1 and Hamamatsu NanoZoomer S60, courtesy of the following funding (1S10OD032267-01, to Denise Ramirez) and the Whole Brain Imaging Core at UTSW. Small animal imaging was provided by the UTSW Pre-Clinical MRI Core supported by the Cancer Prevention Research Institute of Texas (CPRIT RP210099) and the UTSW Pre-Clinical Radiation Core Facility (PCIRCF). The North Texas Clinical Pharmacology Core is supported by the Cancer Prevention Research Institute of Texas (CPRIT RP210209). T.G.O. was supported by NCI awards U24CA213274 and R01-CA244841-05 and received support as a Duke Science & Technology Scholar.

## Supplementary Figures

**Supplementary Figure S1.**
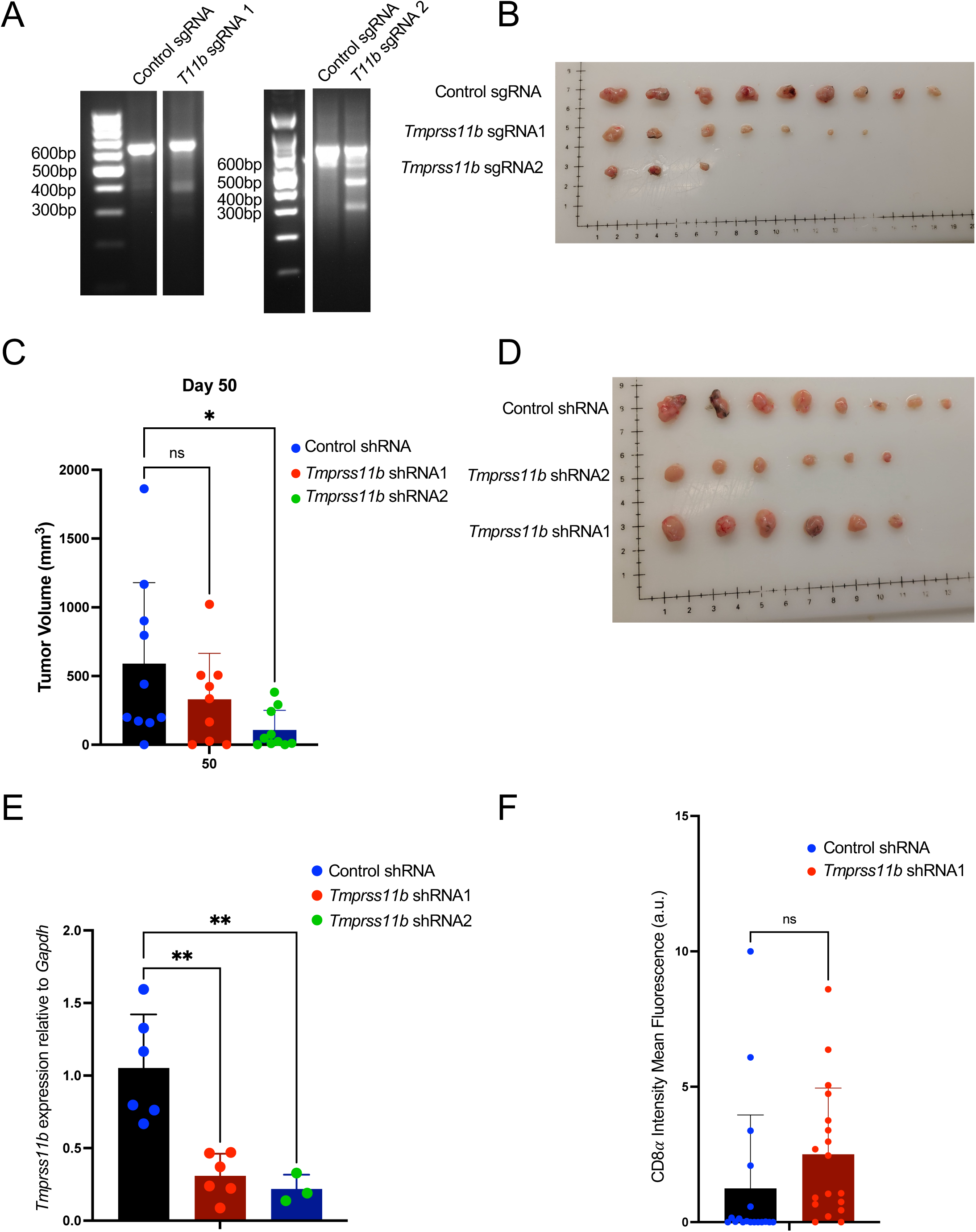
*Tmprss11b* depletion inhibits tumor burden in a syngeneic mouse model of LUSC. **A)** Agarose gel electrophoresis images of PCR amplified products from the Surveyor assay performed on the genomic DNA isolated from KLN205 cells expressing control or *Tmprss11b* sg1 or *Tmprss11b* sg2. **B)** Image showing the resected tumors at endpoint from the syngeneic experiment in Fig 1C. **C)** Quantification of tumor volumes of KLN205 cells expressing doxycycline-inducible control or *Tmprss11b* shRNA on day 50 (terminal measurement) post injection in syngeneic DBA/2 wild-type mice (n=10 control shRNA mice; n=8 *Tmprss11b* sh1 mice; n=10 *Tmprss11b* sh2 mice). Ordinary one-way ANOVA with Dunnett’s multiple comparisons test was used for the statistical analysis. **D)** Image showing the resected tumors from the syngeneic experiment in Fig 1D. **E)** qRT-PCR analysis of *Tmprss11b* mRNA in the resected KLN205 tumors from D). **F)** Quantification of CD8a staining from fluorescent immunohistochemistry (IHC-F) performed on KLN205 tumor sections. An unpaired t test with Welch’s correction was used for the statistical analysis (n=10-14 fields per tumor section, 3 tumors per group).

**Supplementary Figure S2.**
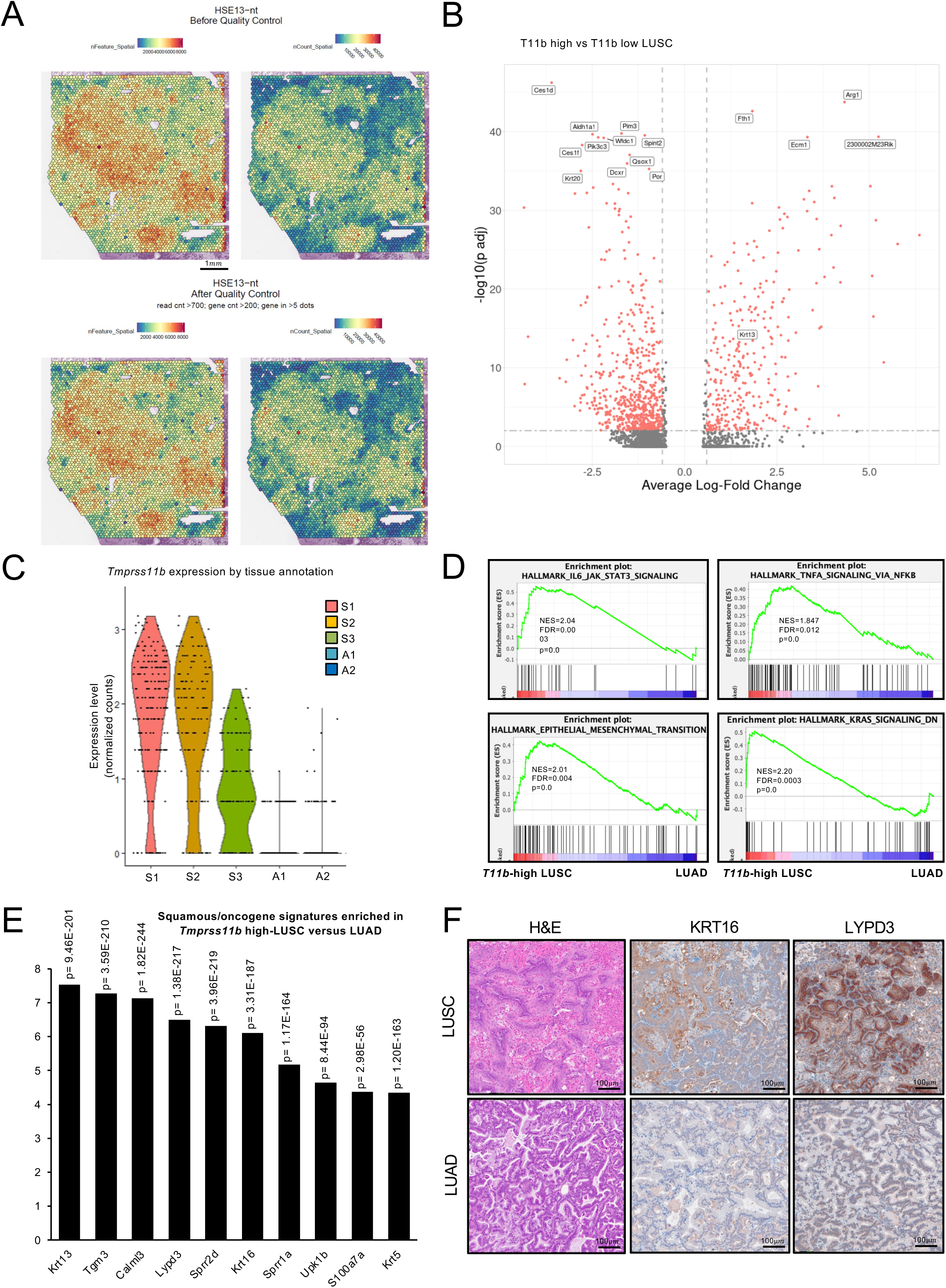
*Tmprss11b*-high squamous tumors have increased expression of oncogenes and squamous markers. **A)** Spatial plots depicting the read counts before and after the quality control process. **B)** Volcano plot showing the top differentially expressed genes in the *Tmprss11b*-high versus low LUSCs spatial data. **C)** Violin plot depicting normalized counts for *Tmprss11b* transcript in the annotated regions (from Fig 2B). **D)** Gene set enrichment analysis (GSEA) of the *Tmprss11b*-high LUSC versus LUAD spatial data with normalized enrichment scores (NES), false discovery rate (FDR) and p values for the indicated gene signatures. **E)** Top squamous markers and known oncogenes from the differential gene expression (DEG) analysis of the *Tmprss11b*-high LUSC versus LUAD spatial data. **F)** Representative H&E and immunohistochemistry (IHC) validation for KRT16 and LYPD3 in LUSC (top) and mucinous LUAD (bottom).

**Supplementary Figure S3.**
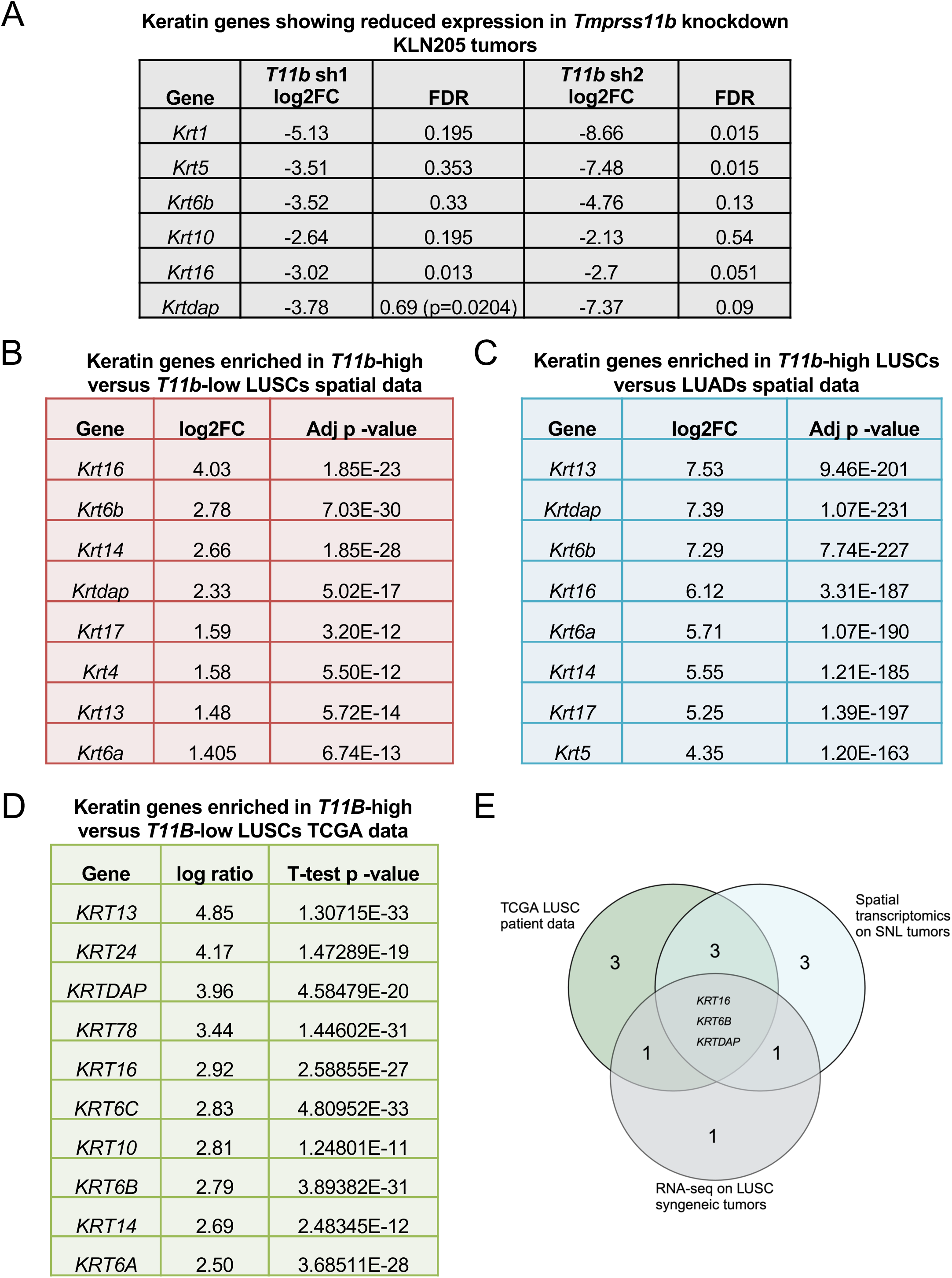
Modulation of TMPRSS11B expression is accompanied by coordinated changes in expression of Keratin genes. **A)** Top downregulated Keratin genes from differential gene expression analysis of control shRNA versus *Tmprss11b* shRNA bulk RNA sequencing from the KLN205 syngeneic experiment in Fig 1D. The log2FC change depicts the reduction in expression of the indicated genes in the *Tmprss11b* knockdown tumors compared to the control. **B)** Top Keratin genes from the differential gene expression (DEG) analysis of the *Tmprss11b*-high versus low in LUSC spatial data from SNL lung tumors. **C)** Top keratin genes from the differential gene expression (DEG) analysis of the *Tmprss11b*-high LUSC versus LUAD spatial data from SNL lung tumors. **D)** Top Keratin genes from the differential gene expression (DEG) analysis of *TMPRSS11B*-high versus low LUSC human tumors from TCGA. **E)** Venn diagram depicting overlapping Keratin genes from the gene lists in A-D.

**Supplementary Figure S4.**
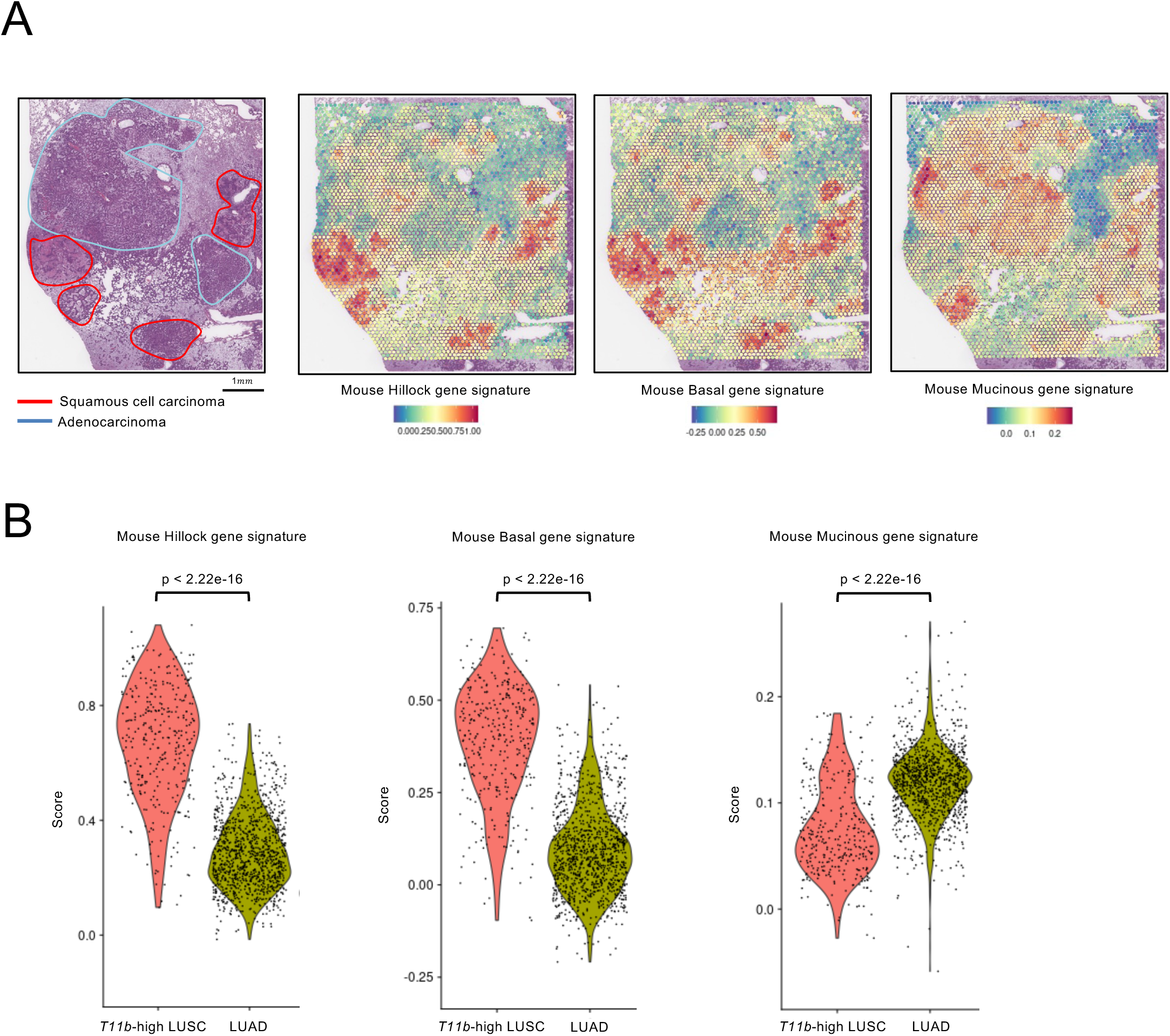
*Tmprss11b*-high squamous tumors show enrichment for hillock (luminal and basal) gene signatures. **A)** H&E image of the SNL lung section used for spatial transcriptomics (left) and spatial plots from the transcriptomic data depicting the distribution of the indicated gene signatures (right). **B)** Violin plots representing the enrichment for the indicated gene signatures in *Tmprss11b*-high LUSC and LUAD.

**Supplementary Figure S5.**
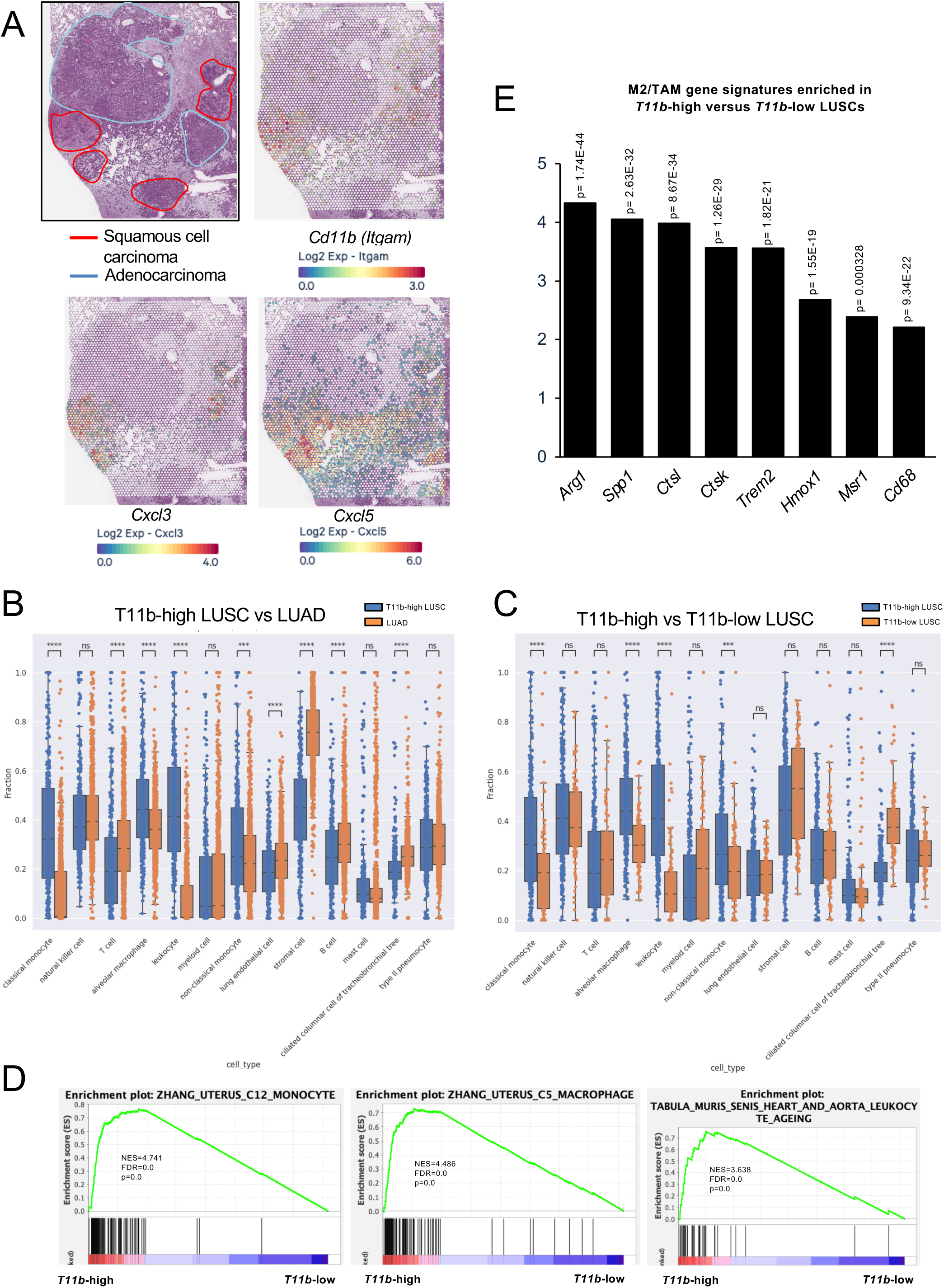
*Tmprss11b*-high squamous tumors show enrichment for M2-like macrophage and neutrophil markers. **A)** H&E image annotated with regions of LUSC and LUAD, and corresponding spatial plots depicting the distribution of the indicated mRNAs (neutrophil markers). **B-C)** Quantification of additional immune cell populations in *Tmprss11b*-high LUSC vs. LUAD (B) and *Tmprss11b*-high vs. Tmprss11b low LUSC (C) using cell deconvolution analysis of the spatial data. **D)** Gene set enrichment analysis (GSEA) of the *Tmprss11b*-high versus low LUSC spatial data with normalized enrichment scores (NES), false discovery rate (FDR) and p values for the indicated immune cell gene signatures. **E)** Bar graph representing top M2-like/TAM genes from the differential gene expression (DEG) analysis of *Tmprss11b*-high versus low LUSC spatial transcriptomics data.

**Supplementary Figure S6.**
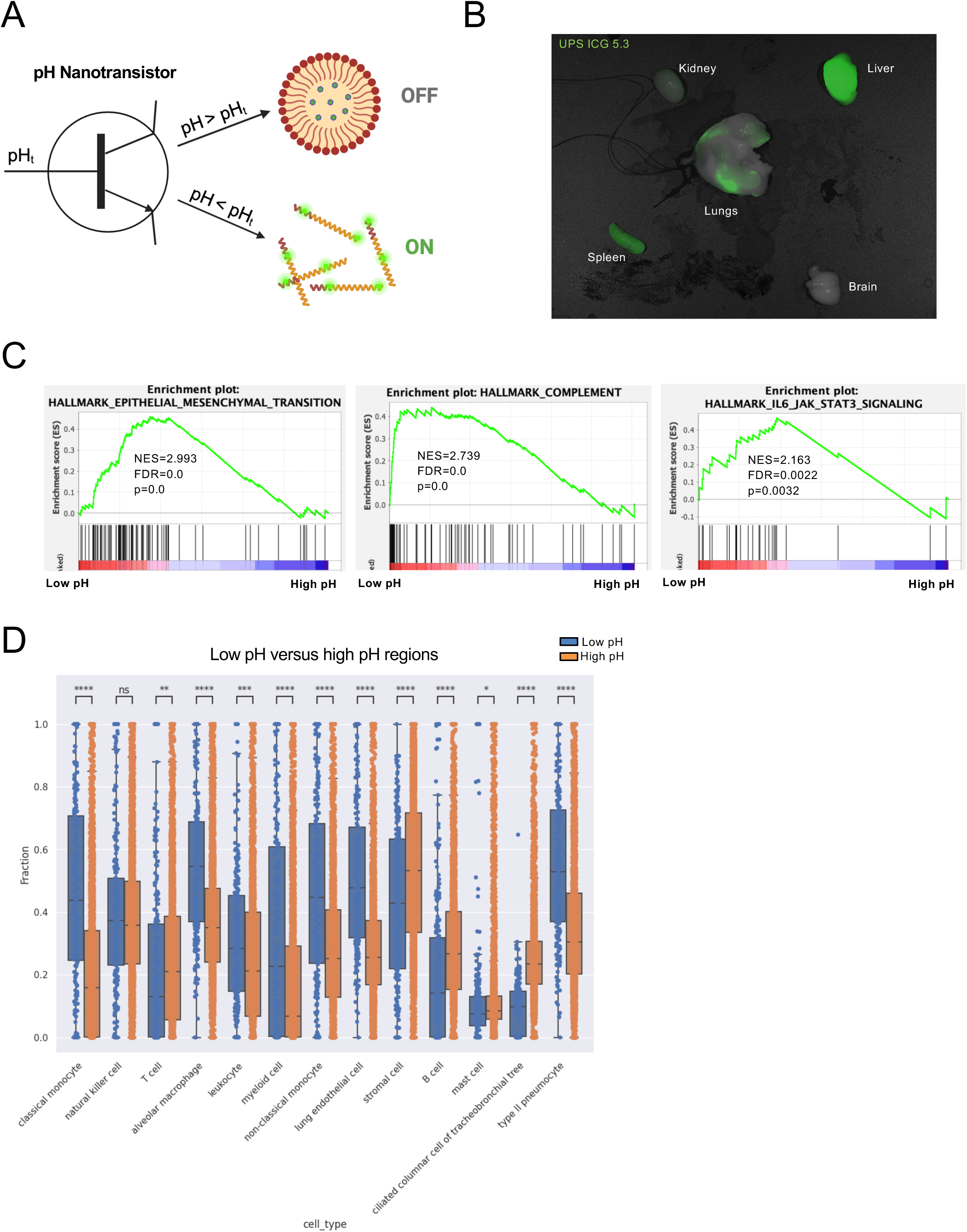
Low pH/acidic regions in the tumor microenvironment show enrichment for oncogenic signaling and immune cell signatures. **A)** Schematic representation of the mechanism of action for the ultra pH sensitive (UPS) nanoparticle, adapted from Feng et al, *Accounts of chemical research*, 2019 (made with BioRender). **B)** Representative fluorescence ICG (UPS 5.3) images of the SNL mouse lung along with liver, kidney, heart, spleen and brain. **C)** Gene set enrichment analysis (GSEA) of the low pH vs. high pH spatial data with normalized enrichment scores (NES), false discovery rate (FDR) and p values for the indicated immune cell gene signatures. **D)** Quantification of immune cell populations in low pH vs. high pH spatial data using cell deconvolution analysis.

**Supplementary Figure S7.**
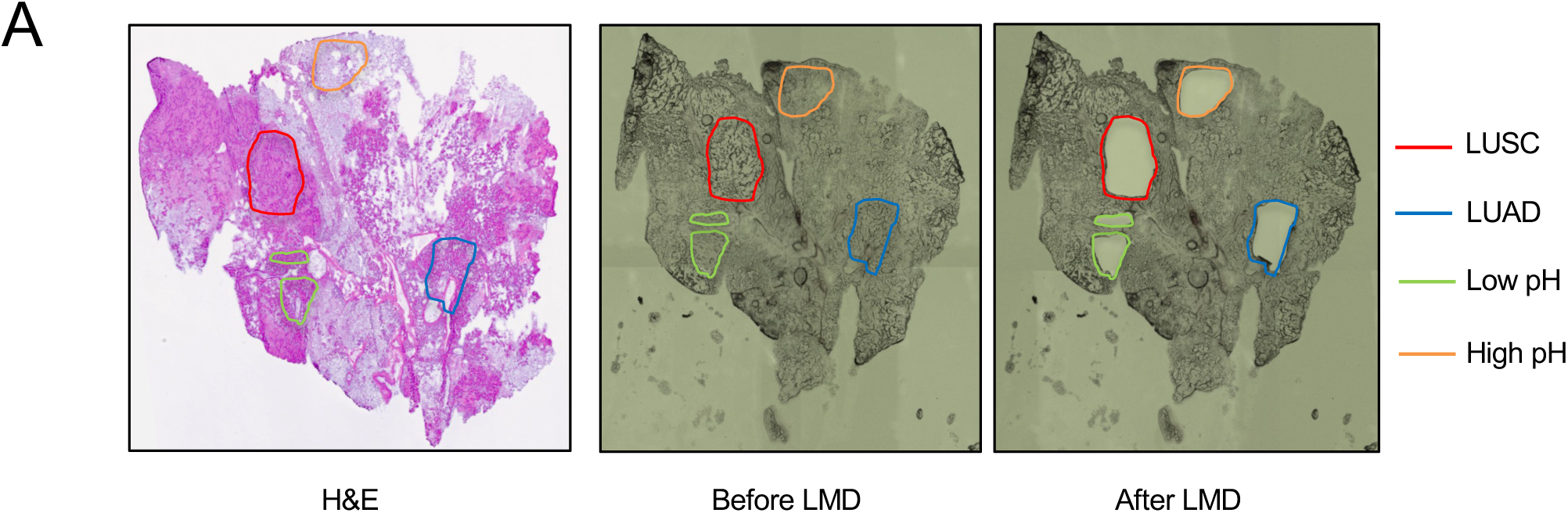
Laser capture microdissection of the regions of interest (LUSC, LUAD, low pH and high pH) using H&E as reference. **A)** Left, Representative H&E image of the SNL lung section (8 months post infection with adeno cre virus) along with corresponding bright field images of the serial section used for LMD, before and after the laser mediated dissection, with annotations.

## Notes

### Competing Interest Statement

KAO serves on the scientific advisory board (SAB) for the Lung Cancer Research Foundation (LCRF).

## REFERENCES

1. Global Burden of Disease Cancer, C., et al. The Global Burden of Cancer 2013. JAMA Oncol 1, 505–527 (2015). 10.1001/jamaoncol.2015.0735

2 Barta, J. A., Powell, C. A. & Wisnivesky, J. P. Global Epidemiology of Lung Cancer. Ann Glob Health 85 (2019). 10.5334/aogh.2419

3 Travis, W. D. Pathology of lung cancer. Clin Chest Med 32, 669–692 (2011). 10.1016/j.ccm.2011.08.005

4 Herbst, R. S., Morgensztern, D. & Boshoff, C. The biology and management of non-small cell lung cancer. Nature 553, 446–454 (2018). 10.1038/nature25183

5 Haslam, A. & Prasad, V. Estimation of the Percentage of US Patients With Cancer Who Are Eligible for and Respond to Checkpoint Inhibitor Immunotherapy Drugs. JAMA Netw Open 2, e192535 (2019). 10.1001/jamanetworkopen.2019.2535

6 Paik, P. K., Pillai, R. N., Lathan, C. S., Velasco, S. A. & Papadimitrakopoulou, V. New Treatment Options in Advanced Squamous Cell Lung Cancer. Am Soc Clin Oncol Educ Book 39, e198–e206 (2019). 10.1200/EDBK_237829

7 Santos, E. S. & Rodriguez, E. Treatment Considerations for Patients With Advanced Squamous Cell Carcinoma of the Lung. Clin Lung Cancer 23, 457–466 (2022). 10.1016/j.cllc.2022.06.002

8 Satpathy, S. et al. A proteogenomic portrait of lung squamous cell carcinoma. Cell 184, 4348–4371 e4340 (2021). 10.1016/j.cell.2021.07.016

9 Hanna, J. M. & Onaitis, M. W. Cell of origin of lung cancer. J Carcinog 12, 6 (2013). 10.4103/1477-3163.109033

10 Jeong, Y. et al. Role of KEAP1/NRF2 and TP53 Mutations in Lung Squamous Cell Carcinoma Development and Radiation Resistance. Cancer Discov 7, 86–101 (2017). 10.1158/2159-8290.CD-16-0127

11 Xu, X. et al. The cell of origin and subtype of K-Ras-induced lung tumors are modified by Notch and Sox2. Genes & development 28, 1929–1939 (2014).

12 Ferone, G. et al. SOX2 is the determining oncogenic switch in promoting lung squamous cell carcinoma from different cells of origin. Cancer cell 30, 519–532 (2016).

13 Zhang, H. et al. Lkb1 inactivation drives lung cancer lineage switching governed by Polycomb Repressive Complex 2. Nat Commun 8, 14922 (2017). 10.1038/ncomms14922

14 Wang, Y. et al. Dysregulated Tgfbr2/ERK-Smad4/SOX2 Signaling Promotes Lung Squamous Cell Carcinoma Formation. Cancer Res 79, 4466–4479 (2019). 10.1158/0008-5472.CAN-19-0161

15 Ruiz, E. J. et al. LUBAC determines chemotherapy resistance in squamous cell lung cancer. J Exp Med 216, 450–465 (2019). 10.1084/jem.20180742

16 Lin, B. et al. Airway hillocks are injury-resistant reservoirs of unique plastic stem cells. Nature 629, 869–877 (2024). 10.1038/s41586-024-07377-1

17 Montoro, D. T. et al. A revised airway epithelial hierarchy includes CFTR-expressing ionocytes. Nature 560, 319–324 (2018). 10.1038/s41586-018-0393-7

18 Plasschaert, L. W. et al. A single-cell atlas of the airway epithelium reveals the CFTR-rich pulmonary ionocyte. Nature 560, 377–381 (2018).

19 Deprez, M. et al. A single-cell atlas of the human healthy airways. American journal of respiratory and critical care medicine 202, 1636–1645 (2020).

20 Updegraff, B. L. et al. Transmembrane Protease TMPRSS11B Promotes Lung Cancer Growth by Enhancing Lactate Export and Glycolytic Metabolism. Cell Rep 25, 2223–2233 e2226 (2018). 10.1016/j.celrep.2018.10.100

21 Bugge, T. H., Antalis, T. M. & Wu, Q. Type II transmembrane serine proteases. J Biol Chem 284, 23177–23181 (2009). 10.1074/jbc.R109.021006

22 Gao, J. et al. Integrative analysis of complex cancer genomics and clinical profiles using the cBioPortal. Sci Signal 6, pl1 (2013). 10.1126/scisignal.2004088

23 Hanahan, D. & Weinberg, R. A. Hallmarks of cancer: the next generation. Cell 144, 646–674 (2011). 10.1016/j.cell.2011.02.013

24 Faubert, B. et al. Lactate Metabolism in Human Lung Tumors. Cell 171, 358–371 e359 (2017). 10.1016/j.cell.2017.09.019

25 Hong, C. S. et al. MCT1 Modulates Cancer Cell Pyruvate Export and Growth of Tumors that Co-express MCT1 and MCT4. Cell Rep 14, 1590–1601 (2016). 10.1016/j.celrep.2016.01.057

26 Dietl, K. et al. Lactic acid and acidification inhibit TNF secretion and glycolysis of human monocytes. J Immunol 184, 1200–1209 (2010). 10.4049/jimmunol.0902584

27 Quinn, W. J., 3rd et al. Lactate Limits T Cell Proliferation via the NAD(H) Redox State. Cell Rep 33, 108500 (2020). 10.1016/j.celrep.2020.108500

28 Watson, M. J. et al. Metabolic support of tumour-infiltrating regulatory T cells by lactic acid. Nature 591, 645–651 (2021). 10.1038/s41586-020-03045-2

29 Huang, X. et al. Neutrophils in Cancer immunotherapy: friends or foes? Mol Cancer 23, 107 (2024). 10.1186/s12943-024-02004-z

30 Shan, T. et al. M2-TAM subsets altered by lactic acid promote T-cell apoptosis through the PD-L1/PD-1 pathway. Oncol Rep 44, 1885–1894 (2020). 10.3892/or.2020.7767

31 Bohn, T. et al. Tumor immunoevasion via acidosis-dependent induction of regulatory tumor-associated macrophages. Nat Immunol 19, 1319–1329 (2018). 10.1038/s41590-018-0226-8

32 Zhang, L. & Li, S. Lactic acid promotes macrophage polarization through MCT-HIF1alpha signaling in gastric cancer. Exp Cell Res 388, 111846 (2020). 10.1016/j.yexcr.2020.111846

33 Li, Z. et al. Lactate in the tumor microenvironment: A rising star for targeted tumor therapy. Front Nutr 10, 1113739 (2023). 10.3389/fnut.2023.1113739

34 Bonde, A. K., Tischler, V., Kumar, S., Soltermann, A. & Schwendener, R. A. Intratumoral macrophages contribute to epithelial-mesenchymal transition in solid tumors. BMC Cancer 12, 35 (2012). 10.1186/1471-2407-12-35

35 Bahr, J. C., Li, X. Y., Feinberg, T. Y., Jiang, L. & Weiss, S. J. Divergent regulation of basement membrane trafficking by human macrophages and cancer cells. Nat Commun 13, 6409 (2022). 10.1038/s41467-022-34087-x

36 Afik, R. et al. Tumor macrophages are pivotal constructors of tumor collagenous matrix. J Exp Med 213, 2315–2331 (2016). 10.1084/jem.20151193

37 Sangaletti, S. et al. Macrophage-derived SPARC bridges tumor cell-extracellular matrix interactions toward metastasis. Cancer Res 68, 9050–9059 (2008). 10.1158/0008-5472.CAN-08-1327

38 Maller, O. et al. Tumour-associated macrophages drive stromal cell-dependent collagen crosslinking and stiffening to promote breast cancer aggression. Nat Mater 20, 548–559 (2021). 10.1038/s41563-020-00849-5

39 Sumitomo, R. et al. M2 tumor-associated macrophages promote tumor progression in non-small-cell lung cancer. Exp Ther Med 18, 4490–4498 (2019). 10.3892/etm.2019.8068

40 Ma, J. et al. The M1 form of tumor-associated macrophages in non-small cell lung cancer is positively associated with survival time. BMC Cancer 10, 112 (2010). 10.1186/1471-2407-10-112

41 Hirayama, S. et al. Prognostic impact of CD204-positive macrophages in lung squamous cell carcinoma: possible contribution of Cd204-positive macrophages to the tumor-promoting microenvironment. J Thorac Oncol 7, 1790–1797 (2012). 10.1097/JTO.0b013e3182745968

42 Jackute, J. et al. Distribution of M1 and M2 macrophages in tumor islets and stroma in relation to prognosis of non-small cell lung cancer. BMC Immunol 19, 3 (2018). 10.1186/s12865-018-0241-4

43 Kaneko, T., LePage, G. A. & Shnitka, T. K. KLN205--a murine lung carcinoma cell line. In Vitro 16, 884–892 (1980). 10.1007/BF02619426

44 Mollaoglu, G. et al. The Lineage-Defining Transcription Factors SOX2 and NKX2-1 Determine Lung Cancer Cell Fate and Shape the Tumor Immune Microenvironment. Immunity 49, 764–779 e769 (2018). 10.1016/j.immuni.2018.09.020

45 Liu, C. et al. S100A7 attenuates immunotherapy by enhancing immunosuppressive tumor microenvironment in lung squamous cell carcinoma. Signal transduction and targeted therapy 7, 368 (2022).

46 Kwon, J. et al. USP13 drives lung squamous cell carcinoma by switching lung club cell lineage plasticity. Molecular Cancer 22, 204 (2023).

47 Arora, R. et al. Spatial transcriptomics reveals distinct and conserved tumor core and edge architectures that predict survival and targeted therapy response. Nature Communications 14, 5029 (2023).

48 Hewitt, R. J. & Lloyd, C. M. Regulation of immune responses by the airway epithelial cell landscape. Nature Reviews Immunology 21, 347–362 (2021).

49 Nakamura, S. et al. Morphologic determinant of tight junctions revealed by claudin-3 structures. Nature communications 10, 816 (2019).

50 Kumar, V. et al. A keratin scaffold regulates epidermal barrier formation, mitochondrial lipid composition, and activity. Journal of Cell Biology 211, 1057–1075 (2015).

51 Bae, J. et al. Targeting LAG3/GAL-3 to overcome immunosuppression and enhance anti-tumor immune responses in multiple myeloma. Leukemia 36, 138–154 (2022).

52 Dragomir, A.-C. D., Sun, R., Choi, H., Laskin, J. D. & Laskin, D. L. Role of galectin-3 in classical and alternative macrophage activation in the liver following acetaminophen intoxication. The Journal of Immunology 189, 5934–5941 (2012).

53 Ng, F. et al. Annexin - 1 - deficient mice exhibit spontaneous airway hyperresponsiveness and exacerbated allergen-specific antibody responses in a mouse model of asthma. Clinical & Experimental Allergy 41, 1793–1803 (2011).

54 Szabo, R. & Bugge, T. H. Membrane-anchored serine proteases as regulators of epithelial function. Biochem Soc Trans 48, 517–528 (2020). 10.1042/BST20190675

55 He, H. et al. Krüppel-like factor 4 promotes esophageal squamous cell carcinoma differentiation by up-regulating keratin 13 expression. Journal of Biological Chemistry 290, 13567–13577 (2015).

56 Riverso, M., Montagnani, V. & Stecca, B. KLF4 is regulated by RAS/RAF/MEK/ERK signaling through E2F1 and promotes melanoma cell growth. Oncogene 36, 3322–3333 (2017).

57 Hu, D. et al. Interplay between arginine methylation and ubiquitylation regulates KLF4-mediated genome stability and carcinogenesis. Nature communications 6, 8419 (2015).

58 Yan, Y. et al. KLF4-mediated suppression of CD44 signaling negatively impacts pancreatic cancer stemness and metastasis. Cancer research 76, 2419–2431 (2016).

59 Ma, R.-Y., Black, A. & Qian, B.-Z. Macrophage diversity in cancer revisited in the era of single-cell omics. Trends in immunology 43, 546–563 (2022).

60 Yu, T. et al. Modulation of M2 macrophage polarization by the crosstalk between Stat6 and Trim24. Nature communications 10, 4353 (2019).

61 Muliaditan, T. et al. Macrophages are exploited from an innate wound healing response to facilitate cancer metastasis. Nature communications 9, 2951 (2018).

62 Yanai, Y. et al. CD8-positive T cells and CD204-positive M2-like macrophages predict postoperative prognosis of very high-risk prostate cancer. Scientific reports 11, 22495 (2021).

63 Hirayama, S. et al. Prognostic impact of CD204-positive macrophages in lung squamous cell carcinoma: possible contribution of Cd204-positive macrophages to the tumor-promoting microenvironment. Journal of thoracic oncology 7, 1790–1797 (2012).

64 Katzenelenbogen, Y. et al. Coupled scRNA-seq and intracellular protein activity reveal an immunosuppressive role of TREM2 in cancer. Cell 182, 872–885. e819 (2020).

65 Wang, C. et al. SPP1 represents a therapeutic target that promotes the progression of oesophageal squamous cell carcinoma by driving M2 macrophage infiltration. British Journal of Cancer, 1–13 (2024).

66 Liu, L. et al. Construction of TME and identification of crosstalk between malignant cells and macrophages by SPP1 in hepatocellular carcinoma. Cancer Immunology, Immunotherapy 71, 121–136 (2022).

67 Wu, N. et al. Cathepsin K regulates the tumor growth and metastasis by IL-17/CTSK/EMT axis and mediates M2 macrophage polarization in castration-resistant prostate cancer. Cell death & disease 13, 813 (2022).

68 Feng, Q., Wilhelm, J. & Gao, J. Transistor-like Ultra-pH-Sensitive Polymeric Nanoparticles. Acc Chem Res 52, 1485–1495 (2019). 10.1021/acs.accounts.9b00080

69 Feng, Q. et al. Severely polarized extracellular acidity around tumour cells. Nature Biomedical Engineering 8, 787–799 (2024).

70 Xu, C. et al. Loss of Lkb1 and Pten leads to lung squamous cell carcinoma with elevated PD-L1 expression. Cancer Cell 25, 590–604 (2014). 10.1016/j.ccr.2014.03.033

71 Reck, M. et al. Pembrolizumab versus chemotherapy for PD-L1–positive non– small-cell lung cancer. New England Journal of Medicine 375, 1823–1833 (2016).

72 Paz-Ares, L. et al. Pembrolizumab plus chemotherapy for squamous non–small-cell lung cancer. New England Journal of Medicine 379, 2040–2051 (2018).

73 Szabo, R. & Bugge, T. H. Membrane-anchored serine proteases as regulators of epithelial function. Biochemical Society Transactions 48, 517–528 (2020).

74 Cassetta, L. & Pollard, J. W. Targeting macrophages: therapeutic approaches in cancer. Nature reviews Drug discovery 17, 887–904 (2018).

75 Conway, E. M. et al. Macrophages, inflammation, and lung cancer. American journal of respiratory and critical care medicine 193, 116–130 (2016).

76 Ruffell, B. & Coussens, L. M. Macrophages and therapeutic resistance in cancer. Cancer cell 27, 462–472 (2015).

77 Sumitomo, R. et al. M2-like tumor-associated macrophages promote epithelial– mesenchymal transition through the transforming growth factor β/Smad/zinc finger e-box binding homeobox pathway with increased metastatic potential and tumor cell proliferation in lung squamous cell carcinoma. Cancer Science 114, 4521–4534 (2023).

78 Wu, P. et al. Inverse role of distinct subsets and distribution of macrophage in lung cancer prognosis: a meta-analysis. Oncotarget 7, 40451–40460 (2016). 10.18632/oncotarget.9625

79 Jiang, B., Zhu, S.-J., Xiao, S.-S. & Xue, M. MiR-217 inhibits M2-like macrophage polarization by suppressing secretion of interleukin-6 in ovarian cancer. Inflammation 42, 1517–1529 (2019).

80 Andrews, S. (Cambridge, United Kingdom, 2010).

81 Dobin, A. et al. STAR: ultrafast universal RNA-seq aligner. Bioinformatics 29, 15–21 (2013).

82 Liao, Y., Smyth, G. K. & Shi, W. featureCounts: an efficient general purpose program for assigning sequence reads to genomic features. Bioinformatics 30, 923–930 (2014).

83 Robinson, M. D., McCarthy, D. J. & Smyth, G. K. edgeR: a Bioconductor package for differential expression analysis of digital gene expression data. bioinformatics 26, 139–140 (2010).

84 Feng, Q., Wilhelm, J. & Gao, J. Transistor-like ultra-pH-sensitive polymeric nanoparticles. Accounts of chemical research 52, 1485–1495 (2019).

85 R Core Team, R. (Vienna, Austria, 2020).

86 Hao, Y. et al. Dictionary learning for integrative, multimodal and scalable single-cell analysis. Nature biotechnology 42, 293–304 (2024).

87 Hafemeister, C. & Satija, R. Normalization and variance stabilization of single-cell RNA-seq data using regularized negative binomial regression. Genome biology 20, 296 (2019).

88 Yu, G., Wang, L.-G., Han, Y. & He, Q.-Y. clusterProfiler: an R package for comparing biological themes among gene clusters. Omics: a journal of integrative biology 16, 284–287 (2012).

89. Wu, T., et al. (2021).

90 Ashburner, M. et al. Gene ontology: tool for the unification of biology. Nature genetics 25, 25–29 (2000).

91 Central, G. et al. The Gene Ontology knowledgebase in 2023. Genetics 224 (2023).

92 Kanehisa, M. & Goto, S. KEGG: kyoto encyclopedia of genes and genomes. Nucleic acids research 28, 27–30 (2000).

93 Kanehisa, M., Furumichi, M., Sato, Y., Kawashima, M. & Ishiguro-Watanabe, M. KEGG for taxonomy-based analysis of pathways and genomes. Nucleic acids research 51, D587–D592 (2023).

94 Cable, D. M. et al. Robust decomposition of cell type mixtures in spatial transcriptomics. Nature biotechnology 40, 517–526 (2022).

95 Elosua-Bayes, M., Nieto, P., Mereu, E., Gut, I. & Heyn, H. SPOTlight: seeded NMF regression to deconvolute spatial transcriptomics spots with single-cell transcriptomes. Nucleic acids research 49, e50–e50 (2021).

96 Dong, R. & Yuan, G.-C. SpatialDWLS: accurate deconvolution of spatial transcriptomic data. Genome biology 22, 145 (2021).

97 Kleshchevnikov, V. et al. Cell2location maps fine-grained cell types in spatial transcriptomics. Nature biotechnology 40, 661–671 (2022).

98 Schaum, N. et al. Single-cell transcriptomics of 20 mouse organs creates a Tabula Muris: The Tabula Muris Consortium. Nature 562, 367 (2018).

99 Subramanian, A. et al. Gene set enrichment analysis: A knowledge-based approach for interpreting genome-wide expression profiles. Proceedings of the National Academy of Sciences 102, 15545–15550 (2005). doi:10.1073/pnas.0506580102

100 Mootha, V. K. et al. PGC-1α-responsive genes involved in oxidative phosphorylation are coordinately downregulated in human diabetes. Nature Genetics 34, 267–273 (2003). 10.1038/ng1180

