## Supplementary Tables S1-S4 for "Transmembrane Serine Protease TMPRSS11B promotes an acidified tumor microenvironment and immune suppression in lung squamous cell carcinoma"

**Supplementary Table 1. Antibodies used in this study.**

| Antigen | Source | Tested Species | Catalog # | Application |
|---|---|---|---|---|
| Phospho-p44/42 MAPK (Erk1/2) (Thr202/Tyr204 | Cell Signaling Technology | Mouse | 4370S | IHC-F |
| CD4 | Cell Signaling Technology | Mouse | 25229S | IHC, IHC-F |
| SOX2 | Cell Signaling Technology | Mouse | 23064S | IHC |
| SPP1 (Osteoponitin) | Abcam | Mouse | ab63856 | IHC |
| TREM2 | Invitrogen | Mouse | PA5-119690 | IHC |
| CD204 (MSR1) | Invitrogen | Mouse | MA5-29733 | IHC |
| HMOX1 | Abcam | Mouse | ab189491 | IHC |
| PDL1 | Cell Signaling Technology | Mouse | 13684S | IHC |
| CD8α | Cell Signaling Technology | Mouse | 98941S | IHC, IHC-F |
| FOXP3 | Ebioscience (Invitrogen) | Mouse | 14-5773-82 | IHC |
| IL-2Rα (CD25) | Abcam | Mouse | ab227834 | IHC |
| CD206 (MRC1) | Abcam | Mouse | ab64693 | IHC |
| ARG-1 | Cell Signaling Technology | Mouse | 93668S | IHC |
| KRT16 | Proteintech (Invitrogen) | Mouse | 17265-1-AP | IHC |
| LYPD3 | Invitrogen | Mouse | PA5-48085 | IHC |
| HNF4α | Cell Signaling Technology | Mouse | 3113S | IHC |
| p40 (DeltaNp63) | Abcam | Mouse | ab203826 | IHC |

**Supplementary Table 2. Primers used in this study.**

**Genotyping Primers for SNL GEMM model**

| Primer | Target | Sequence (5'-3') |
|---|---|---|
| WS268 (Fwd) | Sox2 | GTT ATC AGT AAG GGA GCT GCA GTG G |
| WS270 (Rev-Flox) | Sox2 | AAG ACC GCG AAG AGT TTG TCC TC |
| WS271 (Rev-wt) | Sox2 | GGC GGA TCA CAA GCA ATA ATA ACC |
| NKX Fwd | Nkx2-1 | TTT CTC TCT TGC GGG CTC TA |
| NKX Rev | Nkx2-1 | GGA GGG GCG AGT AGA GAG AG |
| 11FWD | Lkb1 | ATC GGA ATG TGA TCC AGC TT |
| 11REV | Lkb1 | ACG TAG GCT GTG CAA CCT CT |

**Primers for Surveyor assay in Tmprss11b knockout cells**

| Primer | Sequence (5'-3') |
|---|---|
| mT11B_Sur_1_FP | AGAGACTCTTGGGGATGCTG |
| mT11B_Sur_1_RP | GTGTGGGATAGGATGGAGGA |
| mT11B_Sur_2_FP | ACTGCTCTTCCAGGGTTCCT |
| mT11B_Sur_2_RP | ACCAGGACTTGTGAGGATGG |
| mT11B_Sur_3_FP | ACTTGCCCTCTTCTGGTGTG |
| mT11B_Sur_3_RP | CAACTGCAGGTGGTTTAGAGTC |
| mT11B_Sur_4_FP | TATGGGGTTGGAGAAATGGA |
| mT11B_Sur_4_RP | AGAGCTGCCTTCCACACAGT |
| mT11B_Sur_5_FP | TGCAATGGGCTATAATGCAA |
| mT11B_Sur_5_RP | CTTGCCCCCAAACTAATGAA |

**Guide RNA sequences for CRISPR/Cas9 mediated knockouts**

| sgRNA | Sequence (5'-3') |
|---|---|
| Tmprss11b sg1 | GAATGGTAAACACTACTGTG |
| Tmprss11b sg2 | TTGAACAGAATGCTGCATGT |
| Tmprss11b sg3 | ACCTACGACAGAATCACGGG |

**shRNA sequences targetting *Tmprss11b***

| shRNA | Sequence (5'-3') |
|---|---|
| Tmprss11b sh1 | CAAGGTACCAAATATCTCG |
| Tmprss11b sh2 | TTCCGGAAGACAAACCCGA |
| Tmprss11b sh3 | TAATCTTCTCAGCATCAAC |

**Sequencing Primers**

| Primer | Sequence (5'-3') |
|---|---|
| CMV-Forward | CGCAAATGGGCGGTAGGCGTG |
| hU6-Forward | GAGGGCCTATTTCCCATGATT |
| WPRE-Reverse | CATAGCGTAAAAGGAGCAACA |
| T7-Forward | TAATACGACTCACTATAGGG |

**Supplementary Table 3**
**Chemicals and Reagents used in this study**

| Reagents | Source | Catalog # |
|---|---|---|
| 2,2,2-Tribromoethanol | Sigma Aldrich | T48402-25G |
| 2-Methyl-2-butanol | Sigma Aldrich | 152463 |
| Puromycin | Gibco | A1113803 |
| Doxycycline | RPI | D43020-250.0 |
| Doxycycline hyclate | Millipore Sigma | D9891-10G |
| RNAseOUT | Millipore Sigma | R2020-250ML |
| Tween 20 | Millipore Sigma | P9416-100ML |
| Hexadimethrine bromide | Millipore Sigma | 107689 |
| Fetal bovine serum (FBS) | Millipore Sigma | F2442 |
| Effectene Transfection Reagent | Qiagen | 301427 |
| RNeasy Mini Kit | Qiagen | 74106 |
| Opal 3-Plex Detection kit | Akoya Biosciences | NEL810001KT |
| RNAscope Multiplex Fluorescent Kit v2 | Advanced Cell Diagnostics | 3231100 |
| Guide-it Mutation Detection kit | Takara | 631448 |
| TaqMan Universal qPCR Master Mix | ThermoFisher | 4304437 |
| **Experimental models** | | |
| Mouse-KLN205 syngeneic LUSC cell line | ATCC | CRL-1453 |
| Rosa26$^{LSL\text{-}SOX2\text{-}IRES\text{-}GFP}$; *Nkx2-1* $^{fl/fl}$;*Lkb1* $^{fl/fl}$ (SNL) model | Oliver lab | |
| DBA/2 wild-type mice | Charles River | |
| **Plasmids** | | |
| pLentiCRISPR V2 GFP | Addgene | 82416 |
| pLentiCRISPR V2 Puro | Addgene | 98290 |
| pLentiCRISPR V2 Cre | Addgene | 82415 |
| psPAX2 | Addgene | 12260 |
| pMD2.G | Addgene | 12259 |
| **Softwares and Algorithms** | | |
| Prism 10 | GraphPad | https://www.graphpad.com/features |
| Fiji (ImageJ) | NIH | https://imagej.net/ |
| Zen Blue | Zeiss | https://www.zeiss.com/microscopy/us/products/software/zeiss-zen.html |
| GSEA | Broad Institute | https://www.gsea-msigdb.org/gsea/index.jsp |
| Phenochart | Akoya Biosciences | https://www.akoyabio.com/support/software/ |

**Supplementary Table 4. List of MRMs, compound dependent MS/MS parameters and retention times.**

| S.No | Compound ID | Q1 mass | Q3 mass | Dwell time (ms) | DP | CE | CXP | RT (min) |
|---|---|---|---|---|---|---|---|---|
| 1 | Glucose | 179.88 | 89.05 | 20 | -86 | -25 | -9 | 1.34 |
| 2 | 13C6-Glucose | 185.1 | 92.1 | 20 | -80 | -11 | -11 | 1.35 |
| 3 | Lactate | 89 | 43 | 20 | -80 | -16 | -20 | 2.1 |
| 4 | 13C3-Lactate | 92 | 45.1 | 20 | -80 | -16 | -20 | 2.1 |
| 5 | Pyruvic Acid | 86.87 | 43.04 | 20 | -34 | -11 | -5 | 2.04 |
| 6 | Pyruvic Acid | 86.87 | 32.03 | 20 | -34 | -13 | -15 | 2.01 |
| 7 | 13C1 Pyruvic Acid | 88.04 | 32.03 | 20 | -30 | -15 | -15 | 2.05 |
| 8 | Glutamic Acid | 145.89 | 102.02 | 20 | -40 | -20 | -6 | 1.29 |
| 9 | Glutamic Acid | 145.89 | 128.05 | 20 | -40 | -15 | -7 | 1.29 |
| 10 | 13C5-Glutamic Acid | 151.01 | 106.05 | 20 | -36 | -20 | -6 | 1.29 |
| 11 | Succinic Acid | 116.79 | 72.99 | 20 | -50 | -16 | -4 | 2.67 |
| 12 | Succinic Acid | 116.79 | 98.97 | 20 | -50 | -16 | -5 | 2.67 |
| 13 | D4-Succinic Acid | 120.82 | 77.01 | 20 | -30 | -17 | -4 | 2.62 |
| 14 | Citric Acid | 191 | 87.1 | 20 | -30 | -22 | -9 | 2.15 |
| 15 | Citric Acid | 191 | 110.8 | 20 | -30 | -19 | -9 | 2.15 |
| 16 | D4-Citric Acid | 195.15 | 113.03 | 20 | -30 | -19 | -6 | 2.13 |
